# Seed Longevity is Controlled by Metacaspases

**DOI:** 10.1101/2023.03.19.533321

**Authors:** Chen Liu, Ioannis H. Hatzianestis, Thorsten Pfirrmann, Salim H. Reza, Elena A. Minina, Ali Moazzami, Simon Stael, Emilio Gutierrez-Beltran, Evgenia Pitsili, Peter Dörmann, Sabine D’ Andrea, Kris Gevaert, Francisco Romero-Campero, Pingtao Ding, Moritz K. Nowack, Frank Van Breusegem, Jonathan D. G. Jones, Peter V Bozhkov, Panagiotis N. Moschou

## Abstract

To survive extreme desiccation, seeds enter dormancy that can last millennia. This dormancy involves the accumulation of protective but structurally disordered storage proteins through unknown adjustments of proteolytic surveillance mechanisms. Mutation of all six types II metacaspases (MCAs)-II in the model plant Arabidopsis revealed their essential role in modulating these proteolytic mechanisms. MCA-II mutant seeds fail to properly target at the endoplasmic reticulum (ER) the AAA ATPase Cell Division Cycle 48 (CDC48) to dispose of misfolded proteins. MCA-IIs cleave a CDC48 adaptor, the ubiquitination regulatory X domain-containing (PUX) responsible for localizing CDC48 to the lipid droplets. When cleaved, CDC48-PUX is inactivated and allows a lipid droplet-to-ER shuttling of CDC48, an important step in the regulation of seeds’ lifespan. In sum, we uncover antagonism between proteolytic pathways bestowing longevity.

**One-Sentence Summary:** Metacaspase proteases confer seed longevity by antagonizing CDC48 activity.

## Introduction

The desiccation-associated hormone abscisic acid (ABA) modulates seed dormancy by repurposing pre-existing stress-related networks into a seed dormancy program. This allows the storage of high levels of protein in the seed that protect and nourish the embryo (*1*). These proteins with a high level of structural disorder (*2*), might be expected to activate the unfolded protein response (UPR) (*3*). The UPR is a proteolytic mechanism engaging the proteasome and also slows down translation to help deplete misfolded or even disordered proteins that stress the ER. Constitutive UPR slows down translation by reducing the protein synthesis rate, which is achieved by the induction of the IRE1-dependent RNA degradation (RIDD) pathway that promiscuously degrades RNAs on ribosomes (*4*). However, seeds can maintain high levels of protein accumulation which means that they somehow must be able to monitor proteolytic mechanisms (e.g., the UPR).

The cysteine proteases metacaspases (MCAs) are present in bacteria and all eukaryotes except animals (*5*). In contrast to the animal-specific caspases, MCAs cleave after arginine (R) or lysine (K) but not aspartate (D) (*5*). Whilst some organisms have single MCA genes (e.g., budding yeast), higher plants have multi-member MCA families. For example, the model plant *Arabidopsis thaliana* (hereafter referred to as “Arabidopsis”) has nine MCAs, classified according to their structure as type I or II (with three and six members, respectively; hereafter “MCA-IIs”; **fig. 1a**). MCA-Is modulate pathogen-induced programmed cell death, vasculature development, and protein aggregate clearing (*6-8*), while the plant-specific MCA-IIs are involved in stress responses, wound-induced damage-associated molecular pattern signalling and developmental cell death events such as clearance of cell corpses (*9-12*).

**Fig. 1.**
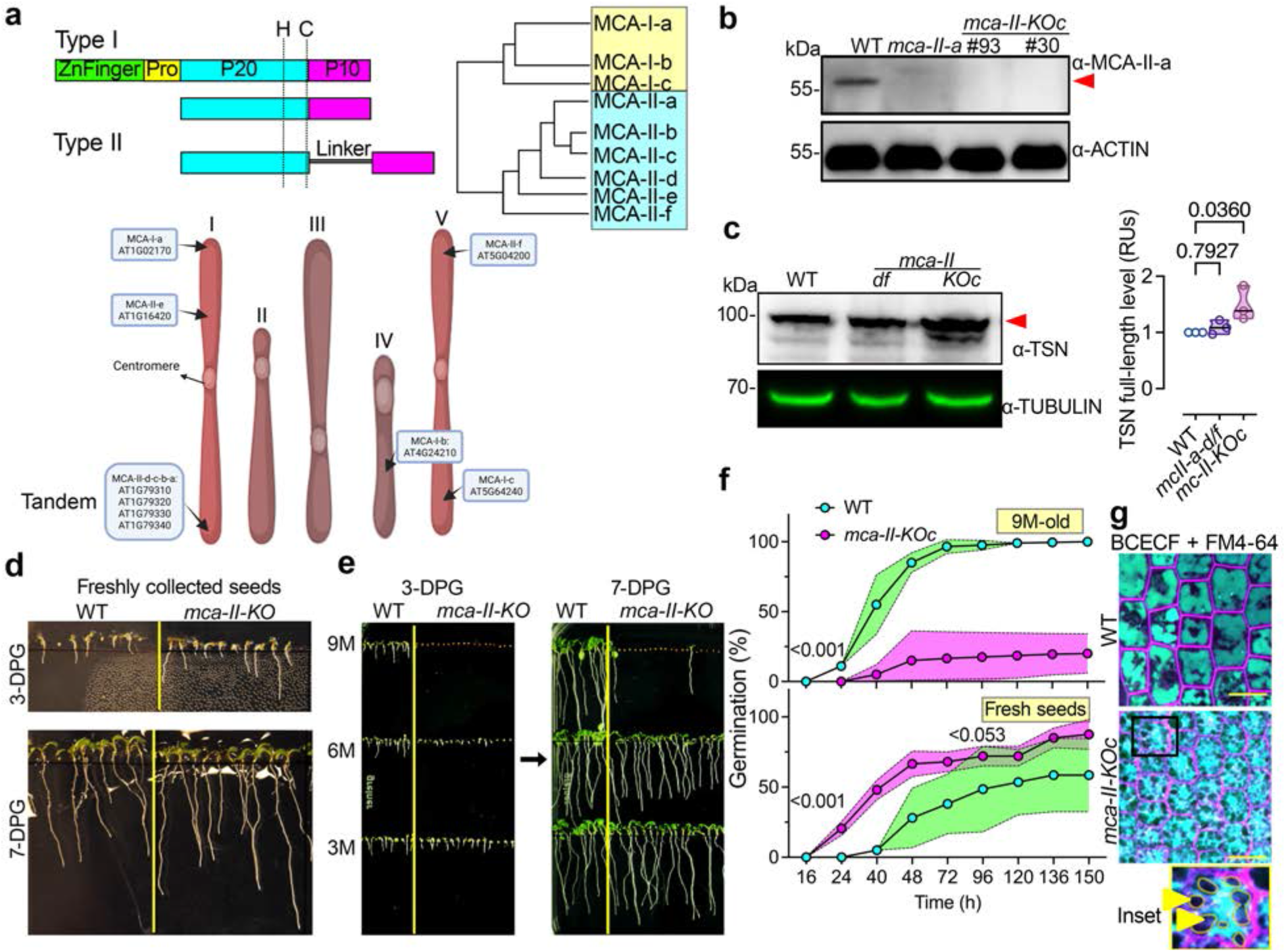
A type II MCA depletion model affects seed physiology and vacuolar morphology. **(A)** Phylogeny of MCAs (type I and II) in Arabidopsis and nomenclature used. The cartoon on the bottom represents all nine MCAs on the five Arabidopsis chromosomes; note the tandem arrangement of four MCA-IIs (MCA-II-a/-b/-c/-d) on chromosome 1. Pro, prodomain; P20 and P10, fragments produced by autoproteolytic cleavage; H, C, catalytic histidine-cysteine pair. **(B)** Representative immunoblot with α-MCA-II-a showing the absence of MCA-II-a in the single *mca-II-a* mutant or the sextuple *mca-II-KOc* (two individual lines #93 and #30), from 7 days post germination (DPG) seedlings. α-ACTIN was used as a loading control and the read arrowhead denotes the signal of MCA-II-a. The experiment was repeated three times (*N* = 3, n = 20 pooled seedlings per lane). **(C)** Representative immunoblot of TSN levels in WT, double *mca-II-df*, or *MCA-II-KOc* mutants from total extracts of 7-DPG. The α-TUBULIN represents loading control. The experiment was repeated three times (*N* = 3, n = 20 pooled seedlings per lane). Right: relative quantification of Tudor Staphylococcal Nuclease (TSN) protein level (red arrowhead, full length) in WT, *mca-II-df* (background used to generate CRISPR mutants), or *mca-II-KOc*. The data are from three experiments and the indicated *P* values were calculated by one-way ANOVA (*N* = 3, n = 3). **(D)** Representative image showing growth of freshly collected seeds of WT and *mca-II-KOc* 3- and 7-DPG. The experiment was repeated three times (*N* = 3, *n* ≥ 40). **(E)** Representative image of WT (9-month-old [9M] seeds) and *mca-II-KOc* (3M-/6M-/9M-old seeds) after 3- and 7-DPG. The experiment was repeated three times (*N* = 3, *n* ≥ 40). **(F)** Germination rate between WT and *mca-II-KOc* seeds (from **d** and **e**). The data are from three experiments, and the indicated *P* values were calculated by one-way ANOVA (*N* = 3, n ≥ 278); the magenta and light green filled areas around the individual points represent ±s.d. **(G)** Vacuole visualization of embryonic roots in WT and *mca-II-KOc* after a 2-day stratification (seeds imbibition) counter-stained with the pH-sensitive lumen dye BCECF. The styryl dye FM4-64 was used for visualization of the plasma membrane (cell contours, magenta). Yellow arrowheads point to vacuole-free regions in the cytoplasm that compresses the vacuolar membrane in the *mca-II-KOc* (corresponding to lipid droplets, see below). Scale bars, 10 μm.

Despite their importance, the exact molecular functions of MCA-IIs remain elusive. This is due to technical difficulties to create double or multiple transfer (T)-DNA-based knockouts because of the tandem arrangement of four out of six MCA-IIs on the first Arabidopsis chromosome (**fig. 1a**, note the nomenclature used for MCA-IIs as in (*5*)).

### Type II Metacaspases have redundant functions

To overcome the potential redundancies among MCA-IIs, we used a CRISPR approach that delivered individual homozygotes for mutations in all MCA-IIs, combinations of them and two transgene-free sextuple independent lines of *MCA-IIs* referred to hereafter as *mca-II-KOc* (***c*** *for “clean line”* without CRISPR transgenes; **fig. S1a-e**). *mca-II-KOc* mutants are likely loss-of-function as they displayed 1) absence of the MCA-II-a protein for which a specific antibody was immunoreactive in immunoblots (unlike other MCA-IIs), accompanied by reduced RNA levels of *MCA-IIs* implying that indel mutations introduced by CRISPR led to non-sense mediated decay known to occur by frameshifts that induce premature stop codons (*13*), 2) reduced cleavage of the known MCA-II substrate TUDOR STAPHYLOCOCCAL NUCLEASE (*14*), and 3) reduced *in vitro* activity on a fluorogenic substrate that is also cleaved by MCA-IIs (ref. (*9*); **figs. 1b-c** and **S1f**).

Remarkably, the *mca-II-KOc* mutant plants displayed only mild developmental phenotypes with reduced leaf serration (i.e., the formation of teeth at leaf margins), earlier flowering, and earlier leaf senescence (**fig. S2a**). However, they did not show signs of compromised developmental cell death, e.g., in the root cap, but showed compromised cell death when challenged with virulent *Botrytis* and *Pseudomonas* (**fig. S2b-d**). In line with our results, a quadruple MCA-IIs mutant showed a similarly compromised response to *Botrytis* (*15*). Furthermore, we observed that effector-triggered immunity (ETI)-associated cell death was abolished when the *mca-II-KOc* were challenged with the effector AvrRps4, which is recognized by Toll-like, Interleukin-1 receptor, Resistance protein (TIR) intracellular nucleotide-binding, leucine-rich repeat (NLR) immune receptors (*16*). On the other hand, no significant differences in AvrRpt2-induced ETI-associated cell death were observed, which is mediated by coiled-coil (CC) NLR receptors. Though it was reported that AvrRpt2-induced ion leakage (i.e., a proxy of the hypersensitive response) was slightly reduced at early time points no significant difference was observed after one day in *mca-II-a* single mutants compared to the WT (*17*). We thus postulate that MCA-IIs play redundant and specific roles in TIR-NLR-mediated cell death response but likely not in CC-NLR-mediated cell death, at least with the effectors used. On the contrary, MCA-IIs may not be involved in the execution of generic cell death programs during development suggesting other modes of action.

During our studies, we observed that freshly collected *mca-II-KOc* seeds germinated faster than WT, suggesting a reduced seed dormancy. When the same batch of seeds was stored for 3 months or more at 4°C as cold can prolong seed lifespan, even under these conditions their germination rate rapidly declined (**fig. 1d-f**). Germination failure was accompanied by fragmented and irregularly shaped vacuoles accumulating aggregates. This vacuolar fragmentation correlated with unknown large structures that appeared to mechanically compress and deform the vacuolar membrane (**fig. 1g**). We turn to this point later, identifying the nature of these structures. The seed germination phenotype was specific for *mca-II-KOc* but was barely observed for the corresponding lower-order mutants. Even the presence of a single active MCA-II, not-targeted by CRISPR could suppress this phenotype (**fig. S3a, b**; note the expression of MCA-IIs in seeds). These results suggest that MCA-IIs appear to redundantly control aspects of seed physiology.

### Metacaspases participate in the ER stress response

As we observed vacuoles that resembled those rich in protein aggregates (storage vacuoles) reported in (ref. (*18*)), we assumed that MCA-IIs may regulate aspects of protein homeostasis in seeds. We postulated that this function could be executed through MCA-IIs proteolytic activity. To identify proteolytic targets of MCA-IIs we first conducted a proteomic analysis of *mca-II-KOc* and WT dry seeds (stored for 3 months at 4°C before seeds of *mca-II-KOc* deteriorate). We used as protein-hit enrichment criterion the semi-quantitation of *mca-II-KOc*/WT peptides with log2FC (fold-change)>1 (**Data S1**). Gene Ontology (GO) term analyses of the enriched proteins showed that *mca-II-KOc* seeds accumulated proteins residing at the endoplasmic reticulum (ER) or associated structures (i.e., lipid droplets), and proteins related to UPR, e.g., the major ER stress-associated chaperones, the binding immunoglobulin protein (BiP) and disulphide isomerases (PDI) (**fig 2a** and **Data S1**; ref. (*19, 20*)). By using Green Fluorescent Protein (GFP) translational fusions of MCA-II-a and -d with expression driven by native promoters, we showed that fluorescently labelled MCA-II-a and -d may associate with the ER in embryonic root cells, and this observation was also validated by cell fractionation in which GFP-MCA-II-a could be detected at the ER fraction (**fig. 2b, c**). These findings suggest that MCA-IIs may directly cleave proteins at the ER to regulate their levels.

**Fig. 2.**
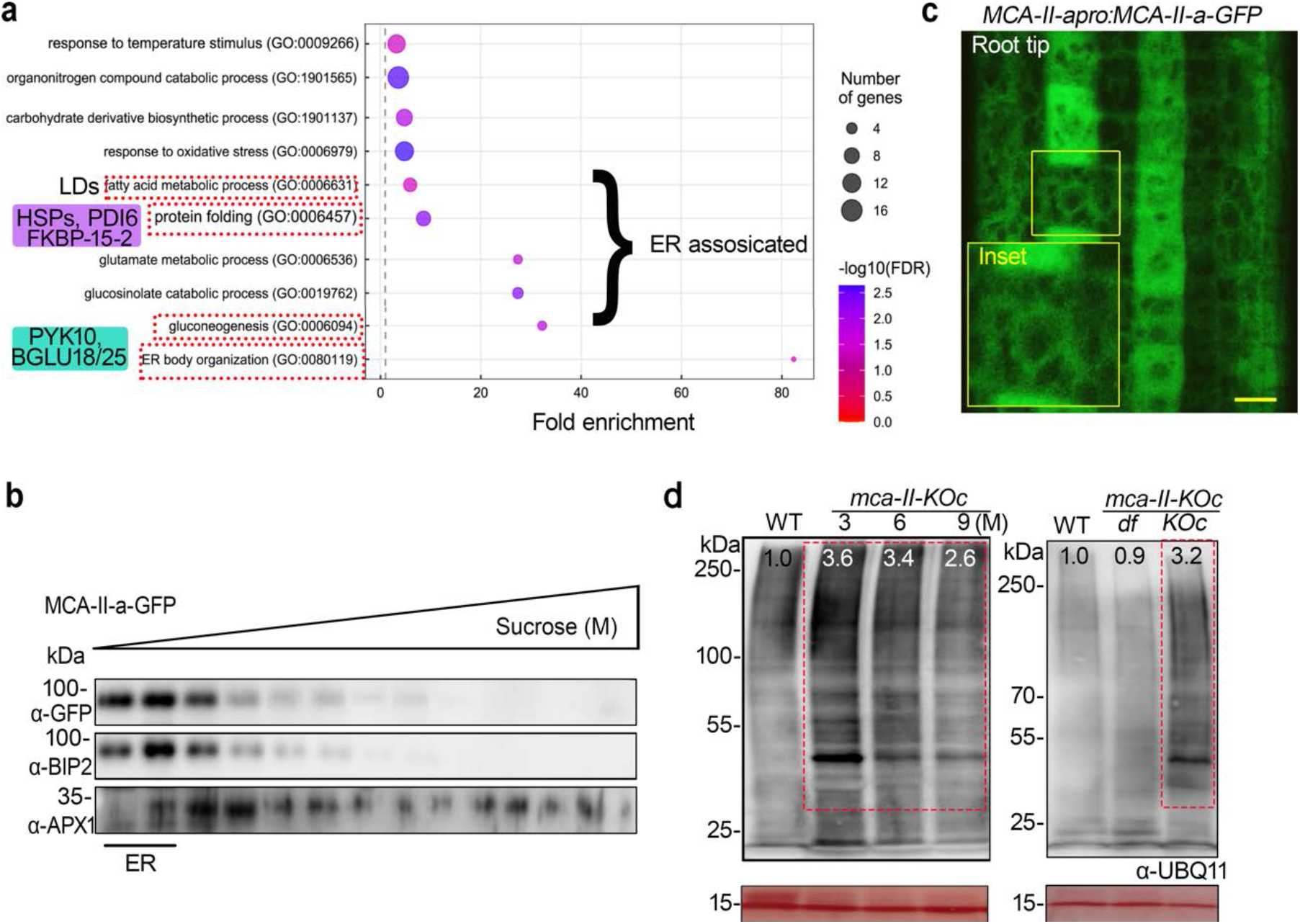
MCA-IIs associate with and affect the ER. **(A)** Enriched GO biological term analysis (log2FC ≥ 1) for total seed proteome of *mca-II-KO*c compared to WT using 3-month-old seeds. The red boxes highlight relevant and overrepresented GO terms, e.g., protein folding and ER body organization (GO:00080119), along with corresponding and abundant genes from the two major terms, e.g., Heat shock 70 kDa proteins (HSP70s) and protein disulfide-isomerase (PDI5, PDI6), Immunoglobulin-binding protein 1 (BIP1: 3.4-fold; BIP2: 2.6-fold; PDI6: 5.03-fold respectively). FKBP15-2 (not found in WT) is involved in ER stress sensing and accelerates the folding of proteins. PYK10 (BGLU23), BGLU18, and BGLU25 are *β*-glucosidases enriched ≥ 100-fold in *mca-II-KOc*. PAG1 (20S proteasome alpha subunit G-1) and PBA1 (20S Proteasome subunit beta 1) were enriched 1.3-fold/not found in WT, which is further verified via immunoblots with α-PAG1 and α-PBA1 in **Fig S5g**. Gluconeogenesis GO term confirms the relevance of this analysis, as MCA-IIs were linked to this process (*41*). The GO term “fatty acid metabolic process” is confirmed later through the findings about oleosin and lipid droplets. FDR, false-discovery rate. **(B)** Representative immunoblot of cell fractionation through sucrose gradient density ultracentrifugation from lines expressing *MCA-II-apro:MCA-II-a-GFP*. BiP2 (ER) and ascorbate peroxidase (APX1) (cytoplasm) were used to assess fraction quality. **(C)** Representative high-resolution confocal micrograph (120 nm axial) from epidermal cells of the meristematic region from embryonic roots upon 2 days imbibition, from lines expressing *MCA-II-apro:MCA-II-a-GFP*. Scale bars, 10 μm. **(D)** Representative immunoblots from seed protein extracts (50 seeds/genotype) with α-UBIQUITIN11(UBQ11). The numbers indicate relative levels compared to Ponceau S staining at the bottom (loading control). Left: WT (9-month-old [9M] seeds) and *mca-II-KOc* seeds harvested at different time points (3-/6-/9 M-old). Right: WT, *mca-II-df, mca-II-KOc* (3 M-old). The experiment was repeated multiple times.

To test whether MCA-IIs directly cleave in vivo the identified proteins accumulating at the ER, we used an N-terminome analysis of the seed proteome by using combined fractional diagonal chromatography (COFRADIC)(*21*). This approach enables the identification of cleavage sites introduced by proteases and the resulting novel N-termini (*neo*-N-termini) formed *in vivo* upon proteolysis (**fig. S3c, d**). COFRADIC analyses, however, did not reveal any of the enriched proteins at the ER as direct targets of MCA-IIs, as these proteins were not enriched in COFRADIC datasets from WT seeds (**fig. S3e** and **Data S1;** i.e.,). Hence, we examined the alternative possibility that *mca-II-KOc* seeds indirectly accumulate the identified ER proteins, by mounting ER stress that is known to activate UPR associated with the accumulation of BiP and PDI (*4*). However, RNA-seq experiments in seeds failed to identify the known transcriptional signatures which associate with UPR in *mca-II-KOc* seeds (**fig. S4a-c** and **Data S2;** ref. (*2*)). Taken together, these results imply that although *mca-II-KOc* seeds show ER stress, they do not activate a constitutive UPR.

Based on these results, we hypothesized that MCAs could function independently from a constitutive UPR. We first tested whether MCA-IIs may regulate the proteasome using MCA-II-a and -b and their inactive **<u>P</u>**roteolytically-**<u>D</u>**ead (MCA-II-a/b^PD^, with catalytic Cys replaced by Ala) variants, as baits in a tandem affinity purification (TAP) approach coupled with liquid-chromatography tandem mass spectrometry (**fig. S5a**). These assays showed that MCA-IIs interacted weakly with two chaperones involved in proteasome assembly (2 out of 32) while mutating the catalytic Cys (MCA-II-a/b^PD^) facilitated these interactions (**fig. S5b** and **Data S3**). The mutant MCA-II-a^PD^ baits but not the WT variants of MCA-II could associate with (poly)ubiquitinated (Ub) proteins, normally recognized by the proteasome, suggesting that MCA-IIs could preferentially target or associate with Ub-proteins (**fig. S5c, d**). Correspondingly, Ub-proteins were enriched in *mca-II-KOc* (**fig. 2d**). Also, proteasome regulatory subunits were enriched in *mca-II-KOc* as revealed by the total proteomic analysis in *mca-II-KOc*, and, in particular, an accumulation of the core 20S proteasome was found, a phenotype reminiscent of proteasomal mutants, e.g., *rpn10* (ref. *(22)*) (**fig. S5e-g**). Furthermore, *mca-II-KOc* mutants were insensitive to proteasomal inhibitor MG132 treatment (a peptide aldehyde that reversibly inhibits the proteasome), but not to other proteolytic inhibitors, as has been reported for several proteasomal mutants (**fig. S5h** ref. (*23*)). Notably, MCA-II-a did not cleave K48 tetra-Ub isopeptide linkages, which represent the most abundant and canonical degradation signals (**fig. S6**; ref. (*24*)). This finding speaks against the possibility that MCA-IIs remove Ub from proteins. Taken together, these results suggest that MCA-IIs may modulate the proteasome.

### Type II metacaspases regulate lipid droplet-associated degradation pathway in seeds

As the UPR was not induced while proteasomal activity was compromised in *mca-II-KOc* mutants, we assumed that Arabidopsis seeds may rely on alternative pathways for protein homeostasis. We focused on ER-associated degradation (ERAD), as it would still promote retrotranslocation of Ub-proteins from ER to the cytosol for degradation by the proteasome much like the UPR (*25*). Furthermore, ERAD fits well in the context of seeds’ proteostasis by allowing excess accumulation of secreted seed storage proteins (e.g., 2S albumins, late embryo abundant, and 12S globulins) that help and protect the embryo, unlike the UPR pathway that would compromise production of secreted proteins by eliciting translational suppression through RIDD (*25, 26*). In plants, ERAD promotes the formation of an impermeable cuticle, a waxy layer attenuating desiccation stress by retaining water in the plant body (*27*). The enzymes for cuticle formation are ER-localized and the cuticle also renders young seedlings (e.g., 3 days old) less responsive to ABA and is required for proper apical hook formation (*28*), which is indispensable for protecting the delicate shoot apical meristem. *rpn10* seeds display compromised longevity, suppressed hypocotyl elongation, and hypersensitivity to ABA and ER stress (*29*). Accordingly, and in line with the possible link of MCA-II to ERAD, we verified cuticle abnormalities in 3-day-old etiolated seedlings of *mca-II-KOc* with toluidine blue permeability assays (**fig. 3a**, ref. (*28*)). We corroborated these results through the specific staining of cuticles with FY-088 in embryonic roots and through cuticle-absence-related defects, such as sensitivity to ABA and apical hook loss in *mca-II-KOc* (**fig. S7a-c**; ref. (*30*)).

**Fig. 3.**
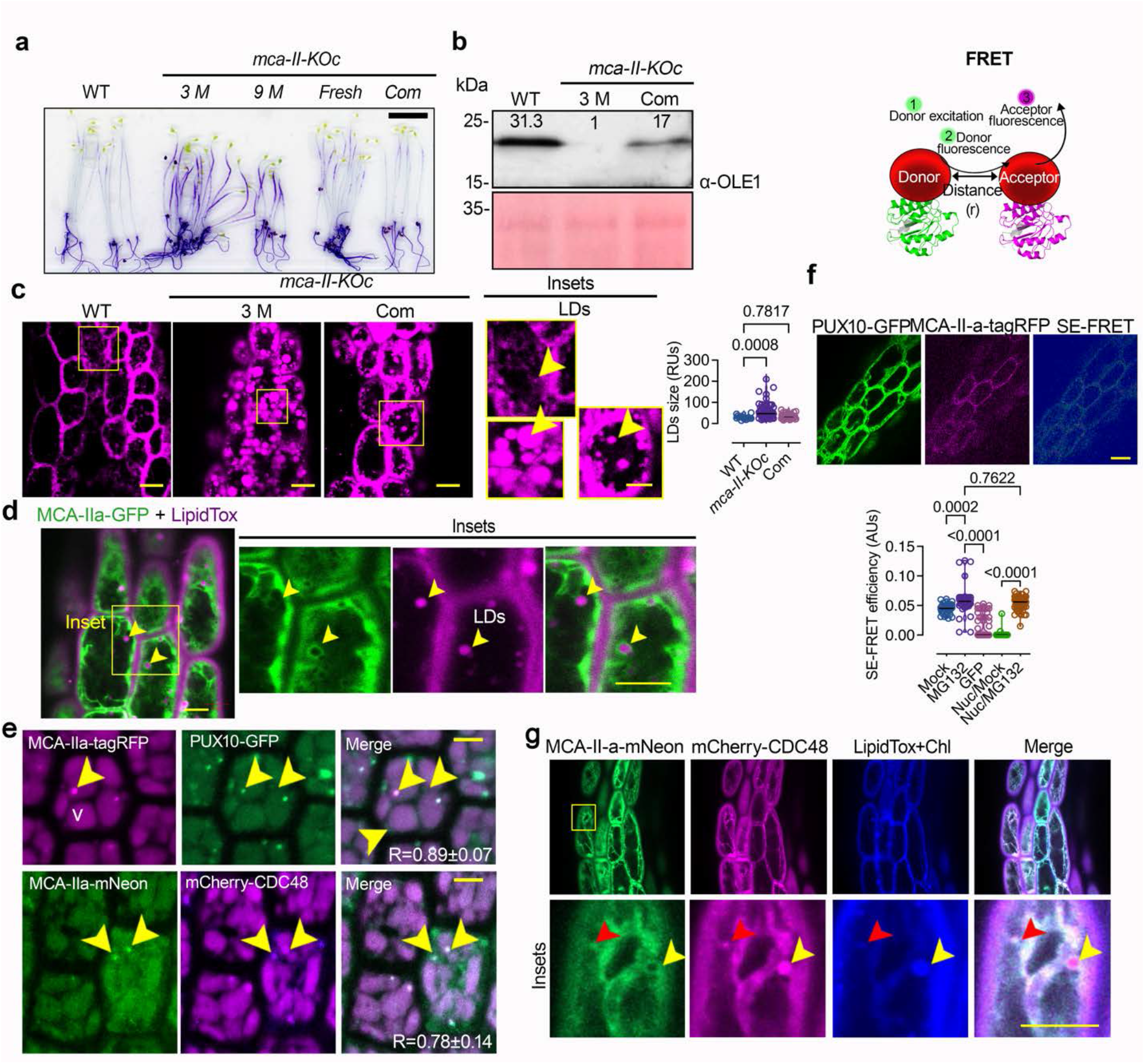
MCA-IIs associate with CDC48/PUX10 in the LDAD pathway. **(A)** Representative images from permeability test of etiolated seedlings (4-days after germination) of WT (9 M-old seeds) and *mca-II-KOc* (FS, fresh seeds/3 M/9 M-old seeds) and complementation line (Com., complemented with *RPS5apro:MCA-II-a-mNeon*; 9 M-old seeds) with toluidine blue (0.05% [w/v]) staining). **(B)** Representative immunoblots of total protein seed extracts of WT, *mca-II-KOc*, and the Com line with an α-Oleosin (3 M-old seeds). The numbers indicate relative levels compared to Ponceau S staining at the bottom (loading control). The experiment was repeated three times (*N* = 3, *n* = 1; 50 seeds/lane). **(C)** Representative micrographs of lipid droplets stained with lipidTox in the hypocotyl region of 2 DPG radicles from WT, *mca-II-KOc*, and the Com line. Yellow arrowheads point out lipid droplet in the insets. Scale bar, 10 μm/5 μm (insets). Right: quantification of lipid droplet sizes. The data are from three experiments, and the indicated *P* values were calculated by ordinary one-way ANOVA (*N* = 3, n ≥ 15). **(D)** Representative confocal micrographs from lines expressing *MCA-II-apro:MCA-IIa-GFP* counterstained with lipidTox in the hypocotyl region of 2 DPG seedlings. Scale bar, 5 μm. The experiment was repeated multiple times. **(E)** Representative confocal micrographs from seeds expressing *MCA-II-apro:MCA-IIa-GFP* with *PUX10pro:PUX10-GFP*. Pearson correlation coefficients (R) were used to estimate colocalizations and are shown on the merged micrographs. Scale bar, 5 μm. **(F)** SE-FRET “sensitized emission” FRET approach, where the emission spectrum of the donor (1) overlaps with the excitation spectrum of the acceptor (2) and if the distance (“r”) between the two molecules is sufficient (i.e., connoting association), energy is transferred (3). **(G)** Representative confocal micrographs from embryo cells showing the colocalization between PUX10/CDC48 and MCA-II-a from lines co-expressing *RPS5apro:MCA-II-a-tagRFP* with *PUX10pro:PUX10-GFP* or *RPS5apro:MCA-IIa-mNeon* with *35spro:mCherry-CDC48a* (radicles produced similar results). The arrowheads denote colocalization between fluorescent signals. Scale bar, 5 μm. The experiment was repeated three times (*N* = 3, *n* = 1). V, vacuole (autofluorescence). Pearson correlation coefficients (R) were used to estimate colocalizations and are shown on the merged micrographs. The experiment was repeated multiple times.

AAA ATPase Cell Division Cycle 48 (CDC48, VCP in vertebrates) is a major ERAD component exporting proteins from the ER for proteasome delivery and also regulates the proteasome, functioning as an “unfoldase/segregase” (*26*). Since CDC48 homologs were enriched 4.1-fold in our proteome dataset (**Data S1**), we tested whether MCA-IIs regulate it. Accordingly, in budding yeast, the sole type I MCA (known as “Mca1”) interacts with CDC48 to regulate protein homeostasis (*31*). We focused on the specialized type of ERAD pathway named “lipid droplets-associated degradation” (LDAD) that removes ubiquitinated oleosins from seed lipid droplets, facilitating their breakdown and remobilization during seed germination (*32*). LDAD regulates oleosin levels at the lipid droplets, while in oleosin mutants, lipid droplets fuse with one another to form larger droplets (*33*). MCA-II-a decorated lipid droplets, and since oleosins control lipid droplet size and stability by preventing lipid droplet coalescence, oleosin deficiency is associated with the formation of larger lipid droplets (*32*). In agreement, we observed a reduction of oleosin in *mca-II-KOc*, which was also confirmed by the increased size of lipid droplets (**fig. 3b, c**). This phenotype was not observed in *mca-II-KOc* complemented with MCA-II-a under the meristem-specific promoter RPS5a (Ribosomal protein S5a from locus AT3G11940). However, *mca-II-KOc* depletion did not affect significantly fatty acids (**fig. S7d**), suggesting that LDAD may not significantly contribute to lipid pools.

In LDAD, the CDC48 adaptor ubiquitination regulatory X (UBX) domain-containing 10 (PUX10), one of the 16 adaptors PUX defined by a ubiquitin-like UBX domain, specifically recognizes Ub-oleosin for degradation, and accordingly, oleosin accumulates in *pu×10* mutants (*32*). PUX10 harbours a ubiquitin-associated (UBA) domain at its N-terminus (*34*); UBA increases the stability of proteins by reducing the incorporation to the proteasome and upon its removal from proteins they get degraded (*35*). We identified CDC48 as a direct interactor of MCA-II-a and confirmed colocalization and direct interactions revealed with a quantitative *in vivo* proximity ligation assay (PLA) and a Fösters Resonance Energy Transfer-sensitized emission (FRET-SE) in *RPS5apro:MCA-II-a-tagRFP/PUX10pro:PUX10-GFP* or CDC48 co-expressing embryonic roots (*36, 37*) (**fig. 3d-f** and **S8a-c**). From these assays, it was evident that in seeds, MCA-II-a, PUX10 and CDC48 colocalize in cellular condensates with higher concentration than the surrounding cytoplasm (**fig. 3f**, arrowhead). These results suggest that MCA-IIs regulate LDAD through an association with CDC48 and PUX10.

### The antagonism between lipid droplet- and ER-associated degradation pathways defines seed longevity

In our colocalization assays, we observed that PUX10-GFP levels in cells were inversely correlated with the levels of MCA-II-a-tagRFP (**fig. 4a**). Conversely, when GFP was fused N-terminally to PUX10, MCA-II-a-tagRFP had no effect on its levels, suggesting that a PUX10 fragment remains stable. To resolve this conundrum, we used the heterologous transient expression system of *Nicotiana benthamiana* leaves (mesophyll cells). This system revealed specific cleavage of PUX10 by MCA-II-a, leading to the accumulation of an MCA-II-a-interacting N-terminal region of PUX10 (**Fig S8d-f**). Accordingly, in our COFRADIC data, the N-terminus of a PUX homolog is highly enriched in WT but not in *mca-II-KOc* (**Fig. S3f**). Hence, although PUX10 is cleaved and the C-terminal part that interacts with oleosin is degraded, the N-terminal fragment containing the stabilizing UBA remains associated with MCA-IIs.

**Fig. 4.**
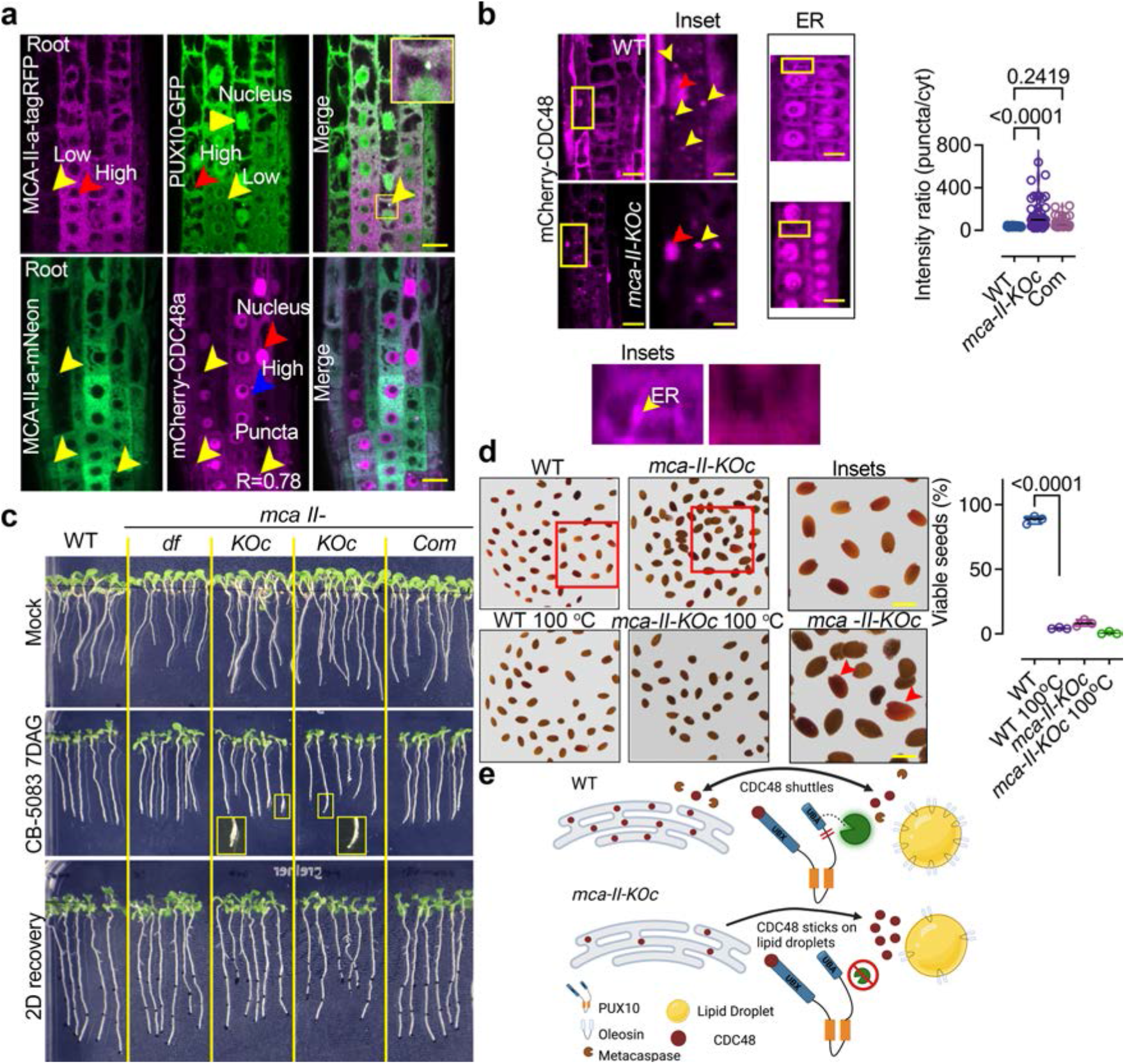
MCA-IIs modulate ERAD by regulating CDC48a localization. **(A)** Representative confocal micrograph of WT expressing *RPS5apro:MCA-II-tagRFP* and *PUX10:PUX10-GFP* in embryonic roots. The yellow arrowheads denote structures reminiscent of lipid droplets, while the red arrowheads denote the nucleus. The inset in the merged micrograph shows the colocalization between the two proteins. Lower: Representative confocal micrograph of WT expressing *RPS5apro:MCA-II-mNeon* with *35Spro:PUX10-GFP* in embryonic roots. The yellow arrowheads denote structures reminiscent of LDs, while the red arrowheads the nucleus. Scale bars, 10 μm. **(B)** Representative confocal micrograph of WT and *mca-II-KOc* expressing *35Spro:mCherry-CDC48a* in embryonic roots 2 days post-imbibition. Scale bars, 10 μm. The yellow arrowheads denote structures reminiscent of lipid droplets, while the red arrowheads the nucleus. Right: quantification of the intensity ratio between the mCherry-CDC48 signal at the ER and cytoplasm. The data are from three experiments and the indicated *P* values were calculated by one-way ANOVA (*N* = 3, n = 8 cells/sample). **(C)** Representative image of 7-day-old seedlings grown on mock (DMSO), in the presence of the CDC48 inhibitor (CB-5083, 2 μM), and following a 2-day recovery (9 days-old seedlings) from CB-5083 on DMSO-containing plates. Note the swelling root tip phenotype observed in *mca-II-KOc* which is indicative of the hypersensitivity to CB-5083 (inset). The experiment was repeated three times (*N* = 3, *n* = 8-10 seedlings/genotype). **(D)** Representative images showing seed viability tests with tetrazolium staining in WT or *mca-II-KOc*. Seeds treated at 100°C represent a positive control. Red seeds are highlighted in the insets (red arrowheads). Right: quantification of viable seeds (% red seeds vs. total seeds). The data are from three experiments and the indicated *P* values were calculated by one-way ANOVA (*N*=3, *n* ≥ 176). **(E)** Suggested model of the role of MCA-IIs in CDC48 regulation in the ER and LDs. In the absence of MCA-IIs PUX10 is not activated and CDC48 is found both in the ER and LDs, while when MCA-IIs are present, PUX10 is cleaved releasing CDC48 to localize at the ER.

We assumed that the lack of PUX10 cleavage in *mca-II-KOc* would likely lead to increased retention of CDC48 attached to lipid droplets. We verified this by showing that in contrast to WT, in *mca-II-KOc* mCherry-CDC48 decorated large intracellular structures reminiscent of lipid droplets, whereas its ER signal decreased (**fig. 4b**). We conjectured that the depletion of CDC48 from ER in the *mca-II-KOc* would lead to aggravated ER-stress and cell death in seeds. To test this, we compared the effect of external inhibition of CDC48 on WT and *mca-II-KOc* using CB-5083, which is a specific inhibitor of CDC48 (ref. (*34*)). Indeed, MCA-II deficiency enhanced the seedling response to CB-5083 (**fig. 4c**, note the root swelling), accompanied by increased cell death in seeds (**fig. 4d**). Together, these results suggest that efficient ERAD requires an MCA-IIs-dependent pathway that involves PUX10 cleavage enabling attenuation of LDAD and ER targeting of CDC48 promoting seed longevity (**fig. 4e**, model).

## Conclusion

Here, we uncovered an antagonism between two proteolytic pathways that involve the specific cleavage of a PUX adaptor to modulate the CDC48 intracellular location. Consistent with our findings, the deletion of PUX10 homologs, ASPL or UBX4, causes ERAD impairment and toxicity in mammal cells and budding yeast (*38, 39*). Furthermore, UBXD8 localization, a mammalian CDC48 adaptor, is regulated by a rhomboid pseudoprotease, which is responsible for shuttling CDC48 between ER and lipid droplets, regulating energy squandering (*25, 40*). In non-plant models, suppression of protein anabolism can lead to enhanced longevity; likewise, plant translational paucity through UPR waves may be functionally equivalent but likely would not suit the seed context in which high protein accumulation is required. The new pathway linking MCA-II-dependent PUX cleavage and CDC48 activity appears to have evolved as an elegant solution to this problem by enabling sustained protein production while maintaining protein homeostasis.

## Acknowledgments

We would like to acknowledge Ikuko Hara-Nishimura and Hannele Tuominen for sharing materials with us.

## Funding

EPIC-XS, Horizon 2020 programme of the European Union project number 823839 (KG, PNM) European Research Council (ERC) Starting Grant ‘R-ELEVATION’ grant 101039824 (PD)

Future Leader Fellowship from the Biotechnology and Biological Sciences Research Council (BBSRC) BB/R012172/1 (PD)

European Union and Greek national funds through the Operational Program Competitiveness, Entrepreneurship, and Innovation, T2EΔK-00597 under the call RESEARCH–CREATE– INNOVATE (“BIOME”; PNM)

European Union Horizon 2020 Marie Curie-RISE PANTHEON grant 872969 (PNM)

Hellenic Foundation of Research and Innovation grant 1426 NESTOR-Theodoros Papazoglou-Always Strive for Excellence (PNM)

Hellenic Foundation of Research and Innovation fellowship 5947 (IHH)

The Swedish Research Council (VR) 2019-04250 (PVB, PNM)

Carl Trygger Foundation (CTS) 22:2025 (PVB)

Knut and Alice Wallenberg Foundation 2018.0026 (PVB, PNM)

European Union Marie Skłodowska-Curie Action IF project 656011 (PNM) DESTINY (BOF-UGent) SS, FVB

## Author contributions

Conceptualization: PNM, PVB, JDGJ, CL

Methodology: CL, IHH, TP, SHR, EAM, AM, SS, EGB, PD, SDA, KG, FRC, PD, MN, FVB

Investigation: CL, IHH, PNM

Visualization: CL, IHH, PNM

Funding acquisition: PVB, PNM

Project administration: PNM

Supervision: PNM

Writing – original draft: PNM, CL, IHH, PVB

Writing – review & editing: all authors

### Data and materials availability

All data, code, and materials used in the analysis are available. All data are available in the main text or the supplementary materials.

## Supplementary Materials

Materials and Methods

Figs. S1 to S8

Data S1 to S3

Table S1

