## Supplemental File for "Seed Longevity is Controlled by Metacaspases"

**The PDF file includes**

Materials and Methods
Figs. S1 to S8
References

**Other Supplementary Materials for this manuscript include the following**

Data S1 to S3
Table S1
Statistics File

### Materials and Methods

#### Plant material

All the plant lines used in this study were in the Arabidopsis Columbia-0 (Col-0) ecotype. The following mutants were used *smb* (*1*), *mca-II-d* (SALK_127688) and *mca-II-f* (GABI_540H06). The two *mca* mutants were used as a background for CRISPR. Primers used for genotyping of mutant lines can be found in **Table S1**. The following transgenic lines used in this study were described previously: *PUX10pro:PUX10-GFP* (*2*) and *MCA-II-fpro:MCA-II-f-GFP* (*3*). Seedlings were grown on half-strength Murashige and Skoog (MS) plant agar media under long-day conditions (16h-light/8h-dark, or as indicated) and were harvested, treated, or examined as indicated in the context of each experiment. In all experiments, seedlings, or plants from T1/F1 (co-localization experiments), T2/F2, or T3/4/5 (for physiological experiments) generations were used. Arabidopsis seeds were sterilized and germinated on half-strength MS agar medium under long-day conditions (16 h light/8 h dark). Arabidopsis plants for crosses, phenotyping of the above-ground part, and seed collection were grown on soil in a plant Aralab chamber at 22°C/19° and a light intensity of 150 µmol $m^{-2}$ $s^{-1}$ with 60% relative humidity stabilized by an infrared sensor. The seeds harvested at the same time under the same condition were used for experiments unless otherwise indicated in the text or figure legends. *Nicotiana benthamiana* plants were grown in Aralab or Percival cabinets at 22°C, 16-h-light/8-h-dark cycles, and a light intensity of 150 µmol $m^{-2}$ $s^{-1}$.

#### Drugs and stainings

The stock solutions of 2 mM FM4-64, 10 mM CB-5083 (1-(4-(benzylamino)-7,8-dihydro-5H-pyrano-(4,3-d)-pyrimidin-2-yl)-2-methyl-1H-indole-4-carboxamide), 1 mM BCECF (2',7'-Bis-(2-Carboxyethyl)-5-(and-6)-Carboxyfluorescein, Acetoxymethyl Ester), LipidTox (1:1000 dilution, Thermo, H34477), ER tracker (1:1000 dilution, Thermo, E34251), Lysotracker (1:1000 dilution, Thermo, L7528), 50 mM MG132, 2 mM Concanamycin A (Con A), 33 mM wortmannin (Wm) and 10 mM E64d were dissolved in dimethyl sulfoxide (DMSO), while 1 M dithiothreitol (DTT) was dissolved in water. Propidium iodide (PI) was dissolved in water. These inhibitors, drugs and stains were diluted in a half-strength MS medium with corresponding concentration and duration, and the final DMSO concentration was ≤0.1% (v/v) in all experiments. Vertically grown 4- to 5-day-old Arabidopsis seedlings were incubated in a half-strength liquid MS medium containing the corresponding drugs for each specific time course treatment as indicated. For confocal microscopy of reporters and mutants, fluorescence images were captured on Leica SP8 or Zeiss780 microscopes and were processed with ImageJ (National Institutes of Health). Cell contours were visualized with propidium iodide (PI) (Molecular Probes) or FM4-64. For the FDA-PI viability staining, seedlings were mounted on a glass slide in FDA solution (1 µL dissolved FDA stock solution [2 mg in 1 ml acetone] in 1 ml of 1/2 MS) supplemented with 10 µg/ml PI. For the seed viability test, tetrazolium red assays were done as described previously (*4*). In short, dry seeds from different genotypes were incubated in the dark in an aqueous solution of 1% (w/v) 2,3,5-triphenyl tetrazolium (TZ) at 28°C for 24 to 48 h with or without indicated treatment. Seeds were rinsed in water before imaging. For the cuticle integrity assay, the experiment was done as described previously (*5*). In short, 48h-post germinated seedlings were incubated in Fluorol Yellow (FY)-088 (0.01% [w/v] in lactic acid) for 70°C, 20min incubation, and the seedlings were rinsed in water and checked under Zeiss 780 microscopes with GFP channel setting (Ex: 488-490 nm, Em: 530-550 nm). For studying the permeability with toluidine blue (*6*), 4 to 5-day-old etiolated seedlings grown on ½ MS plate were collected and then incubated in an aqueous solution of 0.05% [w/v] toluidine blue/0.1% [w/w] Tween 20 for 120 sec followed by a quick washing step in $ddH_2O$. Seedlings were observed with Leica DM6000. For lipid droplet staining, lipidTox (Thermo Fisher Scientific; 500-fold dilution) was used as previously described (*2*).

**Plant infection assay**

The *Botrytis cinerea* strain B05.10 was used in the infection experiments and stored as conidia suspension at −80 °C in 40 % (v/v) glycerol. For conidia production culture was grown in Hydroxyapatite (HA) medium (1% (w/v) Malt extract/ 0.4% (w/v) Glucose / 0.4% (w/v) Yeast extract / 1.5% (w/v) Agar, pH 5.5) and incubated at 25 °C for 7 days. 3-4 weeks plants with fully expanded leaves were used for inoculations. Spore suspensions were prepared in Gamborg Minimal medium (3 g Gamborg B5 basal salt mixtures, 1.36 g $KH_2PO_4$, and 9.9 g glucose per liter), collected by scraping the mycelial colony and adjusted to a concentration of $2 \times 10^5$ spores $ml^{-1}$. A droplet of conidia suspension (10 µl) was placed at one different point of the adaxial surface of each leaf. Control plants were sprayed or drenched with sterile tap water. Disease incidence was measured by counting the number of rotting lesions that appeared on each plant and disease severity was calculated with a disease scale with Fiji software.

**Hypersensitive cell death response phenotyping in Arabidopsis**

The desired avirulent effectors AvrRps4 or AvrRpt2 were delivered to the Arabidopsis plants via a *Pseudomonas fluorescens* effector-to-host analyzer strain (EtHAn), with a *P. syringae* pv. *Syringae* 61 *hrp/hrc* cluster (Type III secretion system machinery) stably integrated into the chromosome in Pf0-1 (Thomas et al., 2009). EtHAn:AvrRps4 and EtHAn:AvrRpt2 were grown on selective KB plates for 24 h at 28 °C (*7*). Bacteria were harvested from the plates, resuspended in infiltration buffer (10 mM $MgCl_2$), and the concentration was adjusted to $OD_{600}$=0.2 (108 CFU $ml^{-1}$). The abaxial surfaces of 5-week-old Arabidopsis leaves were hand infiltrated with a 1 ml needleless syringe. Cell death was monitored 24 h after infiltration.

**Protease activity assay**

Protease activities were measured by fluorogenic peptide-based substrate (EGR-AMC: H-Glu-Gly-Arg-7-amino-4-methylcoumarin) as described previously (*8*). Plant (0.01g seeds or equal amount of 7-day-old seedlings) extract from WT and different mutants using reaction buffer: 50 mM HEPES, pH 7.4, 0.1 % (w/v) 3-[(3-cholamido- propyl) dimethylammonio]-1-propanesulfonate (CHAPS), 50 mm $CaCl_2$, 5 mM dithiothreitol (dTT); EGR-AMC were added at 50 µM final. The release of AMC was measured every 2 min at 30 °C with a Microtiter Plate Fluorometer (Microplate reader Fluostar Omega) using an excitation wavelength of 360 nm and an emission wavelength of 460 nm. Data time points were analyzed by the Omega Fluostar software, and activities were expressed in fluorescence units/min/mg or µg of total protein. Protein concentration was determined using the Bradford reagent (Bio-Rad).

**DNA manipulation and production of transgenic lines**

Electrocompetent Agrobacterium (*Agrobacterium tumefaciens*) strain C58C1 $Rif^R$ (pMP90) or GV3101 $Rif^R$ (i.e., a cured nopaline strain commonly used for infiltration) was used for electroporation, *N. benthamiana* infiltration and floral dip transformation in Arabidopsis (*9*). The following constructs used in the study were described previously: *35Spro:mCherry-PUX10*, *35Spro:mCherry-CDC48a* (*2*). Transcriptional and translational reporters and overexpression constructs used in this study were produced through either Gateway (Invitrogen) cloning using pENTR/D and pENTR5' lines or through GOLDENGATE (Addgene) in the following backbones: (i) pGWB505 and pGWB560 (*10*) (ii) pMDC32(*11*) (iii) (iv) pICSL86900 and pICSL86922

(Addgene). The cDNA of *MCA-II-a* was PCR amplified with Phusion™ High-Fidelity DNA Polymerase & dNTP Mix (Thermo Fisher Scientific, F530N) using cDNA from 7-day-old seedlings. *MCA-II-a-PD* was generated with site-directed mutagenesis with pENTR of *MCA-IIa.* Constructs of *MCA-IIa* or *MCA-IIa-PD* were generated by Gateway cloning with pENTR into different destination vectors which have different tags in N- or C-termini. The coding sequence of *CDC48a, PUX10* were PCR amplified with Phusion™ High-Fidelity DNA Polymerase & dNTP Mix (Thermo Fisher Scientific, F530N) using the cDNA from 7-day-old seedling with pENTR™/D-TOPO™ Cloning Kit (Thermo Fisher Scientific, K240020). Primer sequences used for the amplification of promoters and genes are listed in **Table S1**.

**Construction of MCA-II mutants**

The pICSL binary vector series was utilized to generate CRISPR lines in this study. Primer sequences used for the amplification of promoters and genes are listed in Table S3. To generate the Cas9 expression cassettes, the RPS5a and Cas9z coding sequences and the E9 terminator were amplified using primers flanked with BpiI restriction sites associated with Golden Gate compatible overhangs (**Table S1**). Combinations of three Level 0 vectors containing respectively a promoter, a Cas9z coding sequence and a terminator were assembled in Level 1 vector pICH47811 (Position 2, reverse) by the same 'Golden Gate' protocol but using 0.5 µl of BpiI enzyme (10U/µl, ThermoFisher) instead of 0.5 µl of BsaI-HF. To create the *MCA-IIs* deletion mutant, a previously described multiplexed editing approach was used (*12*). The sgRNAs were designed using the CRISPR-P 2.0 (http://crispr.hzau.edu.cn/CRISPR2) (*13*) and CHOP-CHOP (https://chopchop.cbu.uib.no/) (*14*). To generate the sgRNA expression cassettes, DNA fragments containing the classic or the 'EF' backbone with 7, 67 or 192 bp of the U6-26 terminator were amplified using primers flanked with BsaI restriction sites associated with Golden Gate compatible overhangs (Table S4). The amplicons were assembled with the U6-26 promoter (pICSL90002) in Level 1 vector pICH7751 (gRNA-MCA-II-e, Position 3), pICH7761 (gRNA-MCA-II-b, Position 4), pICH7772 (gRNA-MCA-II-a, Position 5) and pICH7781 (gRNA-MCA-II-c, Position 6) by the 'Golden Gate' protocol using the BsaI-HF enzyme. Combinations of three Level 1 vectors containing a red seed coat maker (FAST-Red, pICSL11015, Position2, *OLE1pro:OLE1-RFP*), a Cas9 expression cassette, and four sgRNA expression cassettes were assembled in Level 2 pAGM4723 (without an overdrive) or pICSL4723 (with an overdrive) by the 'Golden Gate' protocol using the BpiI enzyme. All the plasmids were prepared using a ThermoScientific kit on *Escherichia coli* DH10B electrocompetent cells selected with appropriate antibiotics and X-gal. All the plasmid identification numbers refer to the 'addgene database' (www.addgene.org/). We selected red fluorescing seeds and screened the resulting seedlings for mutation and CRISPR clean lines were selected based on the crossing to WT and to get the segregation lines for further screening of non-red seed coat and resequencing.

**RNA extraction, RNA-seq and quantitative RT-PCR analysis**

Total RNA from the seedlings was extracted using RNeasy Plant Mini Kit with DNaseI digestion (QIAGEN). Reverse transcription was carried out with 500 ng of total RNA using the iScript cDNA synthesis kit (Bio-Rad) according to the manufacturer's protocol. Quantitative PCR with gene-specific primers was performed with the SsoAdvanced SYBR Green Supermix (Bio-Rad) on a CFX96 Real-Time PCR detection system (BioRad). Signals were normalized to the reference genes ACTIN7 using the DCT method and the relative expression of a target gene was calculated from the ratio of test samples to WT. Primer sequences used for the amplification of promoters and genes are listed in **Table S1**. For each genotype, two biological replicates were assayed in three qPCR replicates. qRT-PCR primers were designed using QuantPrime (www.quantprime.de)(*15*). For RNA-seq the concentration of RNA was determined by Qubit®

RNA HS Assay Kit (New England BioNordika BioLab, Q32852). All the RNA samples were treated with DNase I (ThermoFisher Scientific, EN0521) and further enriched with NEBNext® Poly(A) mRNA Magnetic Isolation Module (ThermoFisher Scientific, E7490S). The RNA was measured with Qubit® RNA HS Assay Kits again and libraries were prepared with NEBNext® Ultra™ II RNA Library Prep with Sample Purification Beads (Invitrogen Life Technologies (Ambion Applied Biosystem), E7775S) and NEBNext® Multiplex Oligos for Illumina® (Dual Index Primers Set 1) (New England BioNordika BioLab, E7600S). cDNA library quality was monitored with Agilent DNA 7500 Kit (Agilent Technologies Sweden AB, 5067-1506). cDNA libraries were sequenced with a paired-end sequencing strategy to produce 2 × 150-bp reads using Novogen sequencers and 20 million reads per sample (Novogene, England).

**Cell fractionation**

Cell fractionation was done using *MCA-II-apro:MCA-II-a-GFP* lines based on sucrose gradient density ultracentrifugation (*16*). More specifically, leaves (5g) of the above line were ground followed by protein extraction buffer (50 mM Tris HCl pH 8.2, 2 mM EDTA pH 8.0, 1 mM DTT and protease inhibitors cocktail from Sigma-Aldrich at a 1:100 dilution plus 1 mM phenylmethylsulfonyl fluoride [PMSF]). The extract was filtered through miracloth and centrifuged at 5000 x g for 5 min for the removal of organelles and tissue debris, followed by ultracentrifuge at 100,000 x g for 45 min. Samples were ultracentrifuged overnight at 100.000 g. Subcellular fractions were collected in 2 ml Eppendorf and processed for immunolblot with the following antibodies: α-GFP (Santa Cruz Biotechnology), α-BIP2 (Agrisera), and α-APX1 (Agrisera).

**Fatty acid analysis**

The total fatty acid content and composition of mature seeds were determined by direct transmethylation followed by gas chromatography with a flame ionization detection (*17*).

**Immunocytochemistry, PLA, and imaging**

Immunocytochemistry was done as described previously (*18*). The primary antibodies used were goat anti-CDC48a (VCP1) (diluted 1:500, Abcam. 206320) (*19*). In brief, samples were incubated with primary antibody at 4°C overnight and washed three times with PBS-T , and then incubated for 90 min with Alexa Fluor® 488 AffiniPure Donkey α-Goat IgG (H+L) secondary antibody (Jackson ImmunoResearch, 705-545-147) diluted 1:200-250, After washing in PBS-T and incubating with DAPI (1 µg/mL), specimens were mounted in Vectashield (Vector Laboratories) medium and observed within 48 h. PLA immunolocalization was done as described previously (*20*). Primary antibody combinations diluted 1:200 for α-GFP mouse (Sigma-Aldrich, SAB2702197),1:200 for α-FLAG mouse (Sigma-Aldrich, F1804), 1:200 for α-RFP mouse (Agrisera, AS15 3028) and 1:200 for α-GFP rabbit (Millipore, AB10145) were used for overnight incubation at 4°C. Roots were then washed with MT-stabilizing buffer (MTSB: 50 mM PIPES, 5 mM EGTA, 2 mM $MgSO_4$, 0.1% [v/v] Triton X-100) and incubated at 37°C for 3 h either with α-mouse plus and α-rabbit minus for PLA assay (681 Duolink, Sigma-Aldrich). PLA samples were then washed with MTSB and incubated for 3 h at 37°C with ligase solution as described (Pasternak et al, 2018). Roots were then washed 2x with buffer A (Sigma-Aldrich, Duolink) and treated for 4 h at 37°C in a polymerase solution containing fluorescent nucleotides as described (Sigma-Aldrich, Duolink). Samples were then washed 2x with buffer B (Sigma-Aldrich, Duolink), with 1% (v/v) buffer B for another 5 min, and then the specimens were mounted in Vectashield (Vector Laboratories) medium.

##### Quantification of fluorescent intensity

To create the most comparable lines to measure the fluorescence intensity of reporters in multiple mutant backgrounds, we crossed homozygous mutant bearing the marker with either a WT plant (outcross to yield progeny heterozygous for the recessive mutant alleles and the reporter) or crossed to a mutant only plant (backcross to yield progeny homozygous for the recessive mutant alleles and heterozygous for the reporter). Fluorescence was measured as the mean grey value with subtraction of the background. GFP and chloroplast autofluorescence was excited with the 488-nm line of an argon laser, mCherry and RFP with a 561-nm diode laser, YFP with the 514-nm line of an argon laser, and LipidTOX Deep Red with a 633-nm helium/neon laser. Fluorescence emission was detected between 495 and 510 nm for GFP, 522 to 550 nm for YFP, 600 to 625 nm for mCherry and RFP, and 637 to 650 nm for LipidTOX Deep Red. Chloroplast autofluorescence was imaged between 670 and 700 nm. For multilabeling studies, detection was performed in a sequential line-scanning mode. The apparent diameter of LDs observed by CLSM was estimated using Fiji software (https://fiji.sc/) by manually drawing the diameter using the “line” tool and measuring it with the “measure” function of the software. For BiFC excitation wavelengths and emission, filters were 514 nm/band-pass 530–550 nm for YFP, 561 nm/band-pass 600–630 nm for RFP and 488 nm/band-pass 650–710 nm for chloroplast auto-fluorescence. The objective used was an HC PL APO 40x/1,30 oil CS2 with NA=1.3 (Leica SP8 confocal system).

##### Ubiquitin cleavage assay

Recombinant proteins of GST-PROPEP1, MCA-II-a, MCA-II-a-PD, MCA-II-f, MCA-II-f and Usp2-cc were purified as previously described (*21*) with the protocols described (*22*). The deubiquitylating activity of Usp2-cc, MCA-II-a and MCA-II-f was assayed against recombinant human K48-linked tetra-ubiquitin (BostonBiochem, #UC-210B) by incubating the purified proteases with 2 μg substrate at 37°C for 30 min in either MC-II-f reaction buffer (50 mM MES, pH 5.5, 150 mM NaCl, 10% sucrose, 0.1% CHAPS, 10 mM DTT) or MCA-II-a reaction buffer (50 mM Hepes, pH 7.5, 150 mM NaCl, 10% glycerol, 50 mM $CaCl_2$, 10 mM DTT). Reactions were terminated by adding Sodium dodecyl-sulfate polyacrylamide gel electrophoresis (SDS-PAGE) sample buffer with an additional 50 mM EGTA for MCA-II-a reaction buffer, subjected to SDS-PAGE (12% polyacrylamide) and subsequent silver staining (Invitrogen, #LC6070).

##### Immunoblotting

In general, samples were flash-frozen in liquid $N_2$ and kept at –80°C until further processing. The samples were crushed using a liquid $N_2$-cooled mortar and pestle, and the crushed material was transferred to a 1.5-mL or 15-mL tube. Extraction buffer (EB; 50 mM Tris-HCl pH 7.5, 150 mM NaCl, 10% [v/v] glycerol, 2 mM ethylenediamine tetraacetic acid [EDTA], 5 mM dithiothreitol [DTT], 1 mM phenylmethylsulfonyl fluoride [PMSF], Protease Inhibitor Cocktail [Sigma-Aldrich, P9599] and 0.5 % [v/v] IGEPAL CA-630 [Sigma-Aldrich]) was added according to the plant material used. The lysates were pre-cleared by centrifugation at 16,000 g at 4°C for 15 min, and the supernatant was transferred to a 1.5-mL tube. This step was repeated two times and the protein concentration was determined by the RC DC Protein Assay Kit II (Bio-Rad, 5000122). 2X Laemmli buffer was added, and proteins were separated by SDS-PAGE (1.0 mm thick 4 to 12% [w/v] gradient polyacrylamide Criterion Bio-Rad) in 3-(N-Morpholino) propane sulfonic acid (MOPS) buffer (Bio-Rad) at 150 V or native polyacrylamide gel (Bio-Rad, Any kD TGX gels). Subsequently, proteins were transferred onto polyvinylidene fluoride (PVDF; Bio-Rad) membrane with 0.22-μm pore size. The membrane was blocked with 3% (w/v) BSA fraction V (Thermo Fisher Scientific) in phosphate buffered saline-Tween 20 (PBS-T) for 1 h at room temperature (RT), followed by incubation with horseradish peroxidase (HRP)-conjugated primary antibody at RT for

2 h (or primary antibody at RT for 2 h and corresponding secondary antibody at RT for 2 h). The following antibodies were used: rabbit α-OLE1 (anti-rS3) (*23*), rat α-tubulin (Santa Cruz Biotechnology, 1:1000), rabbit α-GFP (Millipore, AB10145, 1:10,000), mouse α-RFP (Agrisera, AS15 3028, 1:5,000), rabbit α-12S (*24*), rabbit α-APX1 (Agrisera, 1:2000), rabbit α-BIP2 (Agrisera, 1:2000), rabbit α-UBQ11 (Agrisera, 1:2000), goat α-CDC48a (VCP1) (diluted 1:2000, Abcam. 206320), rabbit α-PBA1 (Agrisera, 1:2000), rabbit α-PAG1 (Agrisera, 1:2000), rabbit α-BIP (Agrisera, 1:2000), rabbit α-ACTIN (Agrisera, 1:2000), α-mouse (Amersham ECL Mouse IgG, HRP-linked whole Ab [from sheep], NA931, 1:10,000), α-rabbit (Amersham ECL Rabbit IgG, HRP-linked whole Ab [from donkey], NA934, 1:10,000), α-rat (IRDye® 800 CW Goat anti-Rat IgG [H + L], LI-COR, 925-32219, 1:10.000) and α-rabbit (IRDye ® 800 CW Goat anti-Rabbit IgG, LI-COR, 926-3221, 1:10,000). Chemiluminescence was detected with the ECL Prime Western Blotting Detection Reagent (Cytiva, GERPN2232) and SuperSignal™ West Femto Maximum Sensitivity Substrate (Thermo Fisher Scientific, 34094). The bands were visualized using an Odyssey infrared imaging system (LI-COR).

**Total proteome analysis, COFRADIC and TAP**

An equal number of seeds of WT and *mca-II-KO* were used for the analysis of the total proteome using LC-MS/MS. For COFRADIC, a shortened protocol was used as previously described with the following modifications (*3*). To achieve a total protein content of 1 mg, 0.2 g frozen ground tissue was re-suspended in 1 mL of buffer containing 1% (w/v) 3-[(3-cholamidopropyl)-dimethylammonio]-1-propane sulfonate (CHAPS), 0.5% (w/v) deoxycholate, 5 mM ethylenediaminetetraacetic acid, and 10% glycerol in 50 mM HEPES buffer, pH 7.5, further containing the suggested amount of protease inhibitors (one tablet/10 mL buffer) according to the manufacturer's instructions (Roche Applied Science). The sample was centrifuged at 16,000g for 10 min at 4°C, and guanidinium hydrochloride was added to the cleared supernatant to reach a final concentration of 4 M. Protein concentrations were measured with the DC protein assay (Bio-Rad), and protein extracts were further modified for N-terminal COFRADIC analysis as described previously (*25*). Col-0 (WT) primary amines were labelled with the N-hydroxysuccinimide (NHS) ester of 12C4-butyrate and rsw4 with NHS-13C4-butyrate, resulting in a mass difference of approximately 4 Da between light ($^{12}C_4$) and heavy ($^{13}C_4$) labelled peptides. After equal amounts of the labelled proteomes had been mixed, tryptic digestion generated internal, non-N-terminal peptides that were removed by strong cation exchange at a low pH (*25*). Due to the low amount of input material, the COFRADIC protocol was cut short after the first reversed phase-high performance liquid chromatography (RP-HPLC) step and the resulting 15 fractions were subjected immediately for identification by LC-MS/MS. For TAP, four to five weeks-old Arabidopsis transgenic plants expressing MCA-II-a-TAPa, MCA-II-a-PD-TAPa, MCA-II-b-TAPa ,MCA-II-b-PD-TAPa and sGFP-TAPa were harvested (2-4 g, fresh weight) and ground in liquid $N_2$ in 2 volumes of extraction buffer (50 mM Tris–HCl pH 7.5, 150 mM NaCl, 10% glycerol, 0.1% Nonidet P-40 and 1× protease inhibitor cocktail; Sigma-Aldrich, 1:100 dilution). The analyses and further processing were done as previously described (*26*).

**Visualization of networks and analyses**

Cytoscape 3.5.1 was used. Tab-delimited files containing the input data were uploaded. Unless otherwise indicated, the default layout was an edge-weighted spring-embedded layout, with NormSpec used as edge weight. Nodes were manually re-arranged from this layout to increase visibility and highlight specific proximity interactions. The layout was exported as a PDF and eventually converted to a .TIFF file with Lempel–Ziv–Welch (common name LZW) compression.

#### Quantifications and statistics

The numerical data used in this publication are provided as raw .csv files with the corresponding headings. Graphs were generated by GraphPad Prism or R. PCC is calculated via Fiji, coloc2 tool. All statistical data show the mean ± s.d. (box plots) or the distribution of values (violin plots, kernal density) of at least three biologically independent experiments or samples, or as otherwise stated. Individual data points are on the plots. For violin plots, datasets were smoothed using heavy smoothing which gives a better idea of the overall distribution. In captions, *N* denotes biological replicates, and “*n*” technical replicates or population size. Each data set was tested whether it followed normal distribution when $N \geq 3$ by using the Shapiro normality test. The significance threshold was set at $P<0.05$, and the calculated *P* values are shown in the graphs. Details of the statistical tests applied, including the choice of the statistical method, are indicated in the corresponding figure caption. In boxplots or violin plots, upper and lower box boundaries, or lines in the violin plots when visible, represent the first and third quantiles, respectively, horizontal lines mark the median and whiskers mark the highest and lowest values. For Wilcoxon, *P* values are two-tailed, for Kruskal-Wallis *P* values are approximate, for Welch-ANOVA or Welch-Forsythe in multiple comparisons, *P* values are adjusted. For Kolmogorov-Smirnoff, *P* values are approximate. To increase the robustness of analyses when the normality was marginal (judged by the *P*), sometimes more than one test was used as indicated in the caption (data analyses). For acceptor photobleaching to determine the FRET efficiency, FDR corrections were done through the two-stage step-up method of Benjamini, Krieger and Yekutieli.

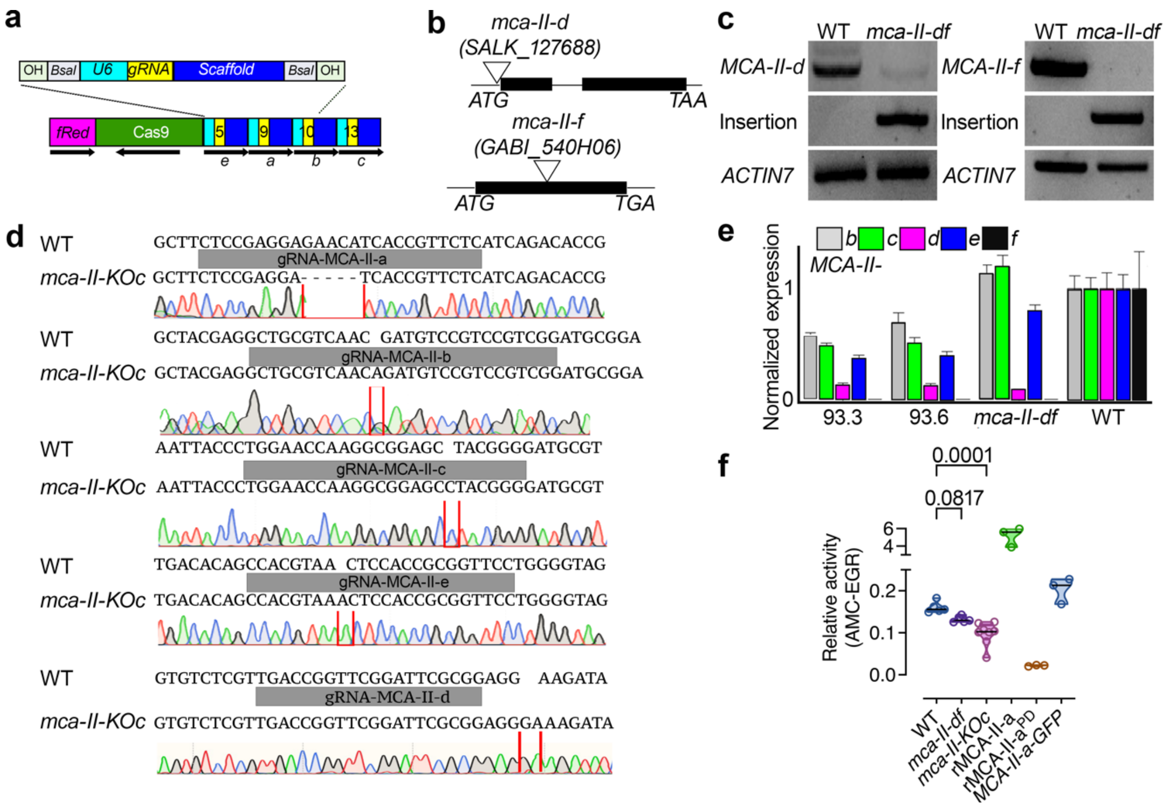


**fig. S1. Generation of a type II MCA depletion model.**

**(A)** Schematic representation of the construct used for the generation of MCA type II sextuple CRISPR mutants. The construct contains a selection marker expressed at the seed coat- (*OLE1pro:Oleosin1-fastRFP; “fRed”*), the Cas9 cassette driven by RPS5apro, and four gRNAs

driven by U6. The gRNAs target the first exon of *MCA-II-a*, *-b*, *-c* and *-e*. Note that Cas and gRNAs are transcribed in opposite directions which we found to increase targeting efficiency. To obtain homozygotes, we screened >1,000 individuals, suggesting a drift from the expected Mendelian segregation.
**(B)** T(transfer)-DNA insert position in the double mutant *mca-II-df* that was used as a background to generate the sextuple CRISPR mutant.
**(C)** RT-PCR from 5-day-old seedlings showing the absence of *MCA-II-d* and *MCA-II-f* transcripts. ACTIN7 was used as a reference gene.
**(D)** Sequencing of the gRNA targeting regions and mutation alignments for *MCA-II-a*, *-b*, *-c*, *-e and -d* in the *mca-II-KOc* mutant line #93.6.
**(E)** Relative expression of *MCA-IIs* (qRT-PCR) after normalization with WT, in *mca-II-df*, and two individual sextuple *mca-II-KOc* mutant lines (#93.3 and 93.6). The data are from two experiments ($N = 2$, $n = 3$).
**(F)** Quantification of the proteolytic activity on the substrate EGR-AMC (H-Glu-Gly-Arg-7-amino-4-methylcoumarin) in WT, *mca-II-df*, and *mca-II-KOc* mutants. As controls, the recombinant purified proteins MCA-II-a (positive) and the proteolytically dead (inactive) variant MCA-II-a$^{PD}$ (negative) were used (proteins purified from *Escherichia coli*, see METHODS), together with the *MCA-IIapro:MCA-II-a-GFP* line. The data are from three experiments, and the indicated *P* values were calculated by one-way ANOVA ($N = 3$, $n = 10$).

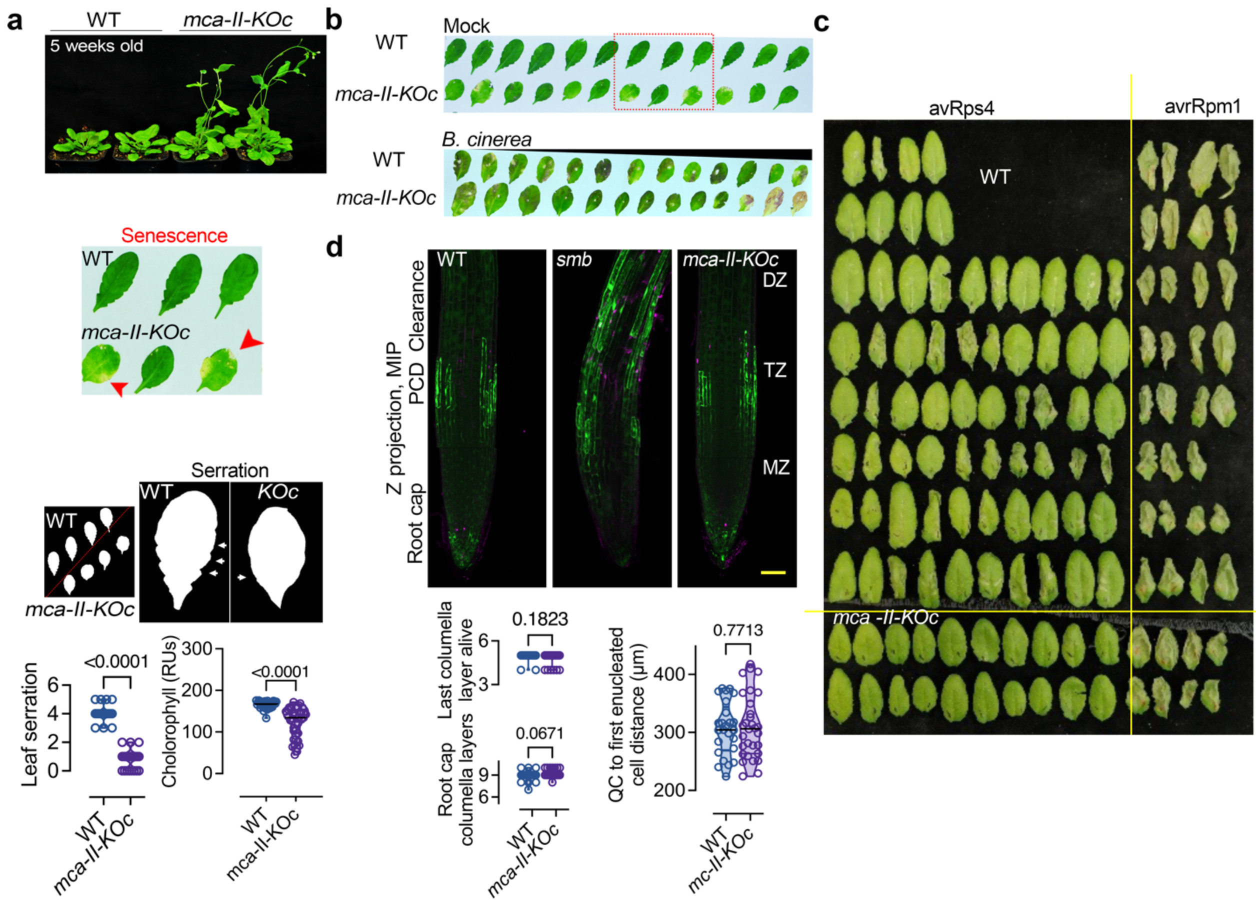


**fig. S2. Phenotypes of a type II MCA depletion model.**

**(A)** Representative image showing the phenotype of 6-week-old WT and *mca-II-KOc* plants. Note the earlier flowering and senescence of *mca-II-KOc* (upper: short-day condition, lower: long-day condition; arrowheads denote leaf yellowing indicative of senescence symptoms). Lower: image showing lack of serration in *mca-II-KOc* leaves, and quantification of chlorophyll and leaf serration in WT and *mca-II-KOc* plants. The data are from three experiments, and the indicated *P* values were calculated by one-way ANOVA ($N = 3, n = 10$).
**(B)** Comparison between WT and *mca-II-KOc* fully expanded leaves with *Botrytis cinerea* infection in short-day conditions.
**(C)** Comparison between WT and *mca-II-KOc* fully expanded leaves with *Pseudomonas* (avRps4 and avrRpm1) infection in short-day conditions. Row 1: wild-type Col-0 (WT); row 2: *rrs1-3 rrs1b-1* double mutant in Col-0; row 3: *mca-II*-d/f double mutant; row 4: 7/8/9 triple mutant; row 5: 5/7/8/9 quadruple mutant; row 6: 6/7/8/9 quadruple mutant; row 7: 4/6/7/8/9 quintuple mutant; row 8: 5/6/7/8/9 quintuple mutant; row 9 and 10: two individual lines of *mca-II-KOc* sextuple mutant. 5-week-old short-day growing plants were infiltrated with the Pf0-1 EtHAn strains, either delivering the effector AvrRps4 or AvrRpm1.
**(D)** Representative confocal micrographs showing root cap programmed cell death assay in WT, *sombrero* (*smb*), and *mca-II-KOc* determined by fluorescein diacetate (FDA; green) and propidium iodide (PI; magenta). FDA stains living cells, while PI dead cells. The experiment was repeated two times ($N = 2$, $n = 20$). The mutant *smb* shows compromised cell death in the root cap and was used as a control here. Scale bar, 20 µm. DZ, differentiation zone; TZ, transition zone; MZ, meristematic zone. Right: quantifications of last columella cell alive, root cap columella

cells, or the distance between the QC (quiescent centre) to the first enucleated cell in the root cap in WT or *mca-II-KOc*. The data are from three experiments, and the indicated *P* values were calculated by one-way ANOVA ($N$ = 3, n ≥ 7).

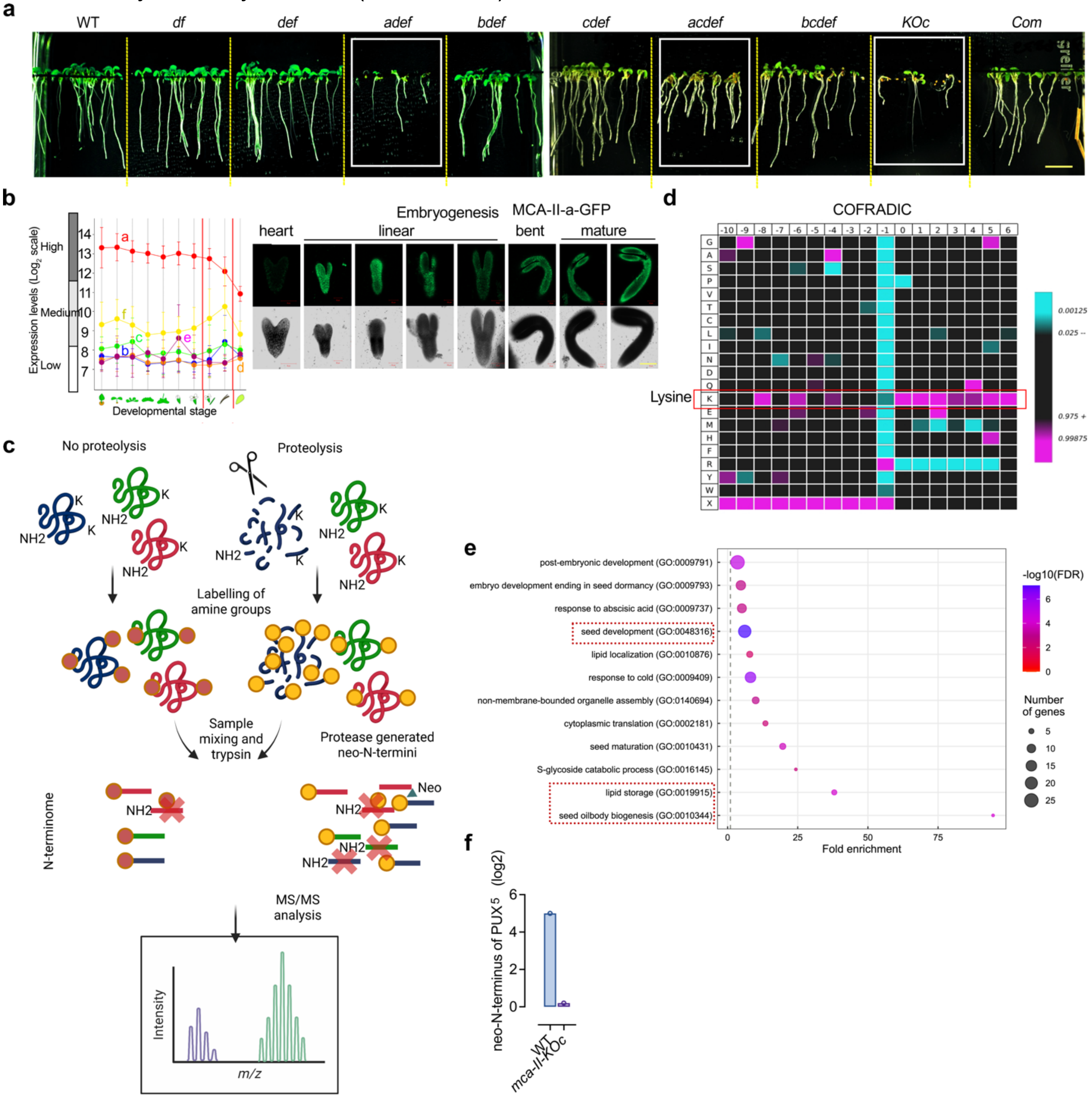


**fig. S3. MCA-IIs redundantly regulate seed physiology and participate in the modulation of LDs.**

**(A)** Seven-day-old Arabidopsis seedlings carrying different combinations of *MCA-II* loss-of-function mutations generated via CRISPR and a complementation line (“Com”) expressing *PRS5apro:MCA-II-a-mNeon* in the *mca-II-KOc* background. Note the correlation between low germination and mutations of MCA-II-a (white frames).

**(B)** Expression profiles of MCA-IIs at different developmental stages determined by Genevestigator (https://genevestigator.com) (left) and representative confocal micrographs of *MCA-II-apro:MCA-II-a-GFP* expression at different stages of embryogenesis (right).

**(C)** Illustration of the workflow of the N-terminomic COFRADIC method (COmbined FRActional DIagonal Chromatography). This method is based on the differential labelling of amine groups of aa between the samples, followed by trypsin digestion and tandem mass spectrometry analysis, detecting the potential endogenous substrates of MCA-IIs using total seeds extract (100 mg, 3M-old, *n* = 5).
**(D)** MCA-IIs cleavage specificity determined using Icelogo analyses from N-termini produced from COFRADIC datasets after sequence alignment and statistical correction for Arabidopsis proteins (pipelines in ref. (*27*)). The heat map shows the aa frequencies at the P6-P10′ positions (P1 is R) and proteins rich in lysines (K) spanning the cleavage site are preferred targets of MCA-IIs at this stage. The data are from two experiments (*N* = 2).
**(E)** GO biological term analyses of the potential endogenous substrates of MCA-IIs ($Log_2FC \geq 1$). Note the enrichment in terms of seed and post-embryonic development, lipid storage, and seed oil body (lipid droplet, LD) biogenesis. The s-glycoside catabolic process corresponds to BGLUs identified in the total proteome (**fig. 2**).
**(F)** Quantification data for PUX10 protein that was highly enriched in WT but not in *mca-II-KOc*.

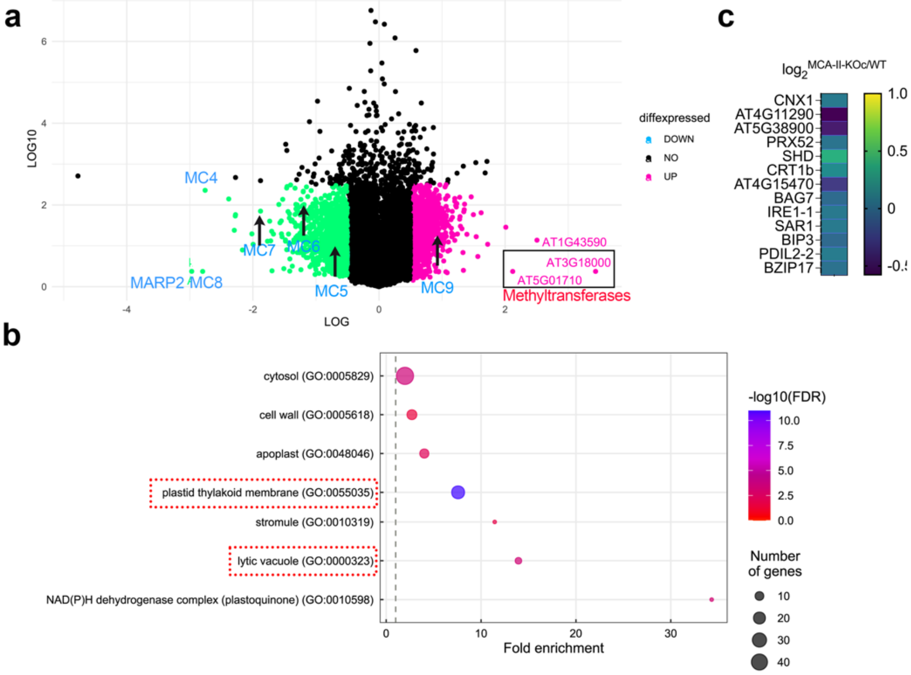


**fig. S4.** ***mca-II-KOc* do not show UPR.**

**(A)** Volcano plot showing differentially expressed genes (DEGs) in *mca-II-KOc* compared to the WT in the RNA-seq dataset obtained from total RNA of 3M-old seeds ($Log_2FC \geq 0.5$ or $\leq -0.5$). The data are from two experiments. Arrows denote the expression levels of MCA-II genes validating the results obtained in the qRT-PCR in **fig. S1**.
**(B)** GO term analysis for Cellular Component (CC) visualization of DEGs ($Log_2FC \geq 0.5$ or $\leq -0.5$) showing a wide range of subcellular localization such as the lytic vacuole or the plastid thylakoid membrane and lumens which is a complex of connected membranes with the lumen of chloroplast where the light reaction of photosynthesis occurs. The enrichment of the lytic vacuole and chloroplast-associated GO terms could be justified due to enriched proteasome complex formation leading to impaired protein homeostasis. A characteristic example is the chloroplast regulation via the proteasome vital for the developmental transitions of plastids to chloroplasts (*28*)

**(C)** Heat map showing that UPR-related marker genes (selected according to ref. (*29*)) were not differentially expressed in our dataset. For example, BIP3 is a chaperone that, along with other proteins, facilitates the proper folding of newly synthesized proteins (*30*). IRE1-1 is protein kinase which is auto-activated in the presence of unfolded proteins in the ER and activates transcription factors such as bZIP60 for UPR initiation (*31, 32*).

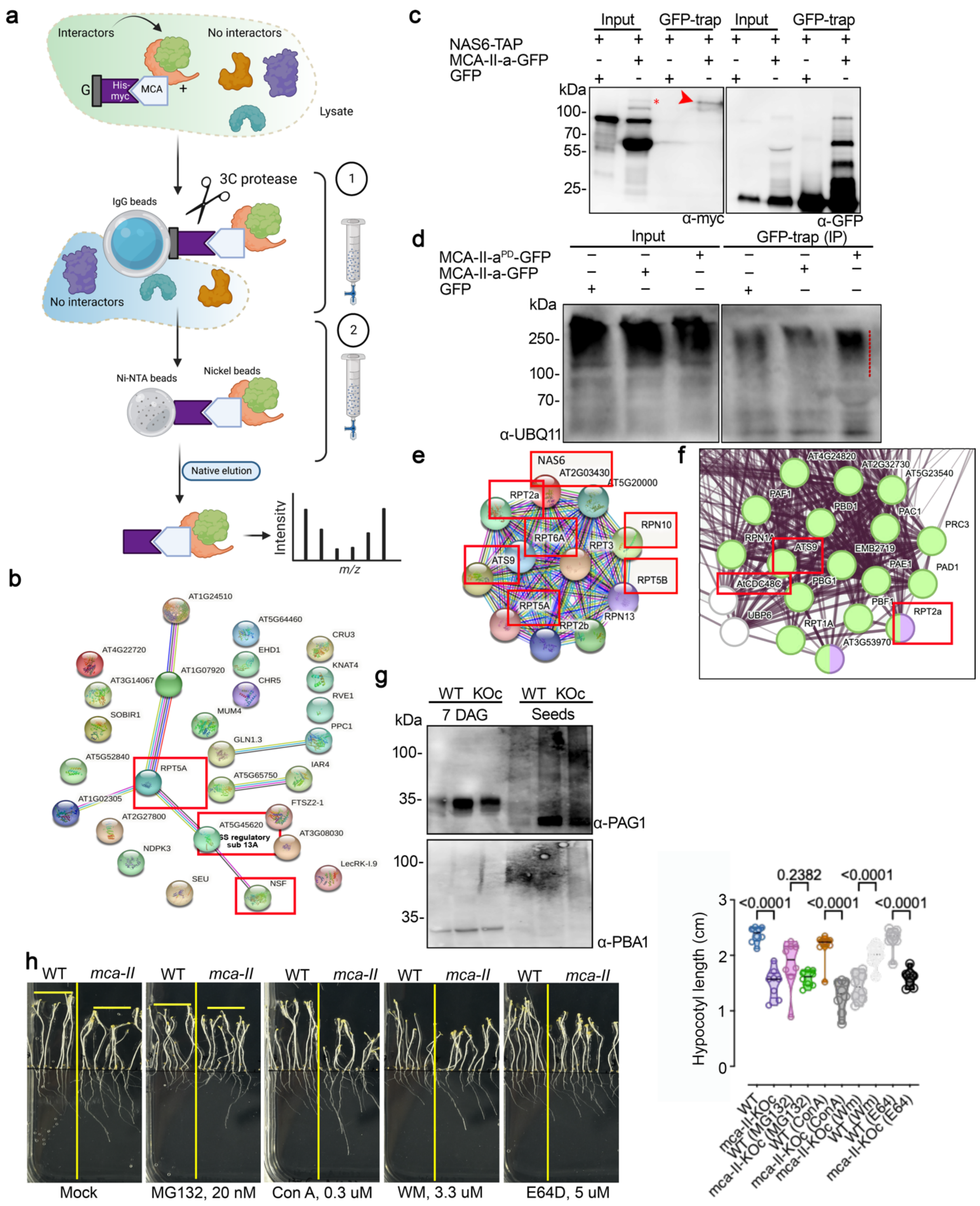

**fig. S5. The interplay of MCA-IIs and proteasome.**

**(A)** Cartoon showing the Tandem affinity purification (TAP) approach with the baits 35spro:MCA-II-a-TAP/35spro:MCA-II-a$^{PD}$ (protease-dead)-TAP, and 35spro:MCA-II-b-TAP/35spro:MCA-IIb$^{PD}$-

TAP using a total extract from 7-day-old transgenic lines to investigate the potential substrates and/or interactors (details in methods). MCA-IIa$^{PD}$ and MCA-IIb$^{PD}$ were used here as a means to capture substrates as the protease-substrate complex may be dissembled after proteolysis (*33*).
**(B)** Proteasomal STRING map showing potential interactors obtained in TAP (red frames) and their relatively weak linkage to the proteasome. Note the red frame marked the network enrichment of proteasome regulatory subunits for RPT5A (26S proteasome AAA-ATPase regulatory subunit particle 5b) and AT5G45620 (proteasome regulatory subunits 13A)
**(C)** Representative immunoblot showing the co-immunoprecipitation of MCA-II-a-GFP (bait) with NAS6-TAP-Myc from total protein extracts from *N. benthamiana* transient expression (4-days-post-infiltration). One red asterisk indicated that the higher order of NAS6 was pulled down with MCA-II-a-GFP. The experiment was repeated two times.
**(D)** Representative immunoblot showing Ub-linked proteins immunoprecipitation with the baits MCA-II-a-GFP or MCA-II-a$^{PD}$-GFP (α-UBQ11). Protein extracts were obtained from 4 days post-infiltration in the *N. benthamiana* transient expression system. Note the red line for poly-Ub-linked protein in the MCA-II-a$^{PD}$-GFP sample. The experiment was repeated two times (N = 2).
**(E)** STRING map showing protein-protein interaction map of RPT5A (Regulatory Particle 5a), one of the six AAA-ATPases of the proteasome regulatory particle) and related proteasome regulatory subunits network including NAS6 (chaperon for proteasome assembly, probable 26S proteasome regulatory subunit p28) which was used in the pull-down studies in **D**. Note the NAS6 and CDC48a are both chaperones for proteasome assembly (*34*). The ATS9 (regulatory particle non-ATPase 6, RPN6) and regulatory particle AAA-ATPase 2A (RPT2a) have been shown in **F** (see below).
**(F)** Part of the STRING map from the total proteome analysis (FC>2 compared to WT) represents a cluster enriched in proteasome complex. The red arrowheads denote CDC48 (FC=4.1, adj P = 0.67). Red cycles represent the proteasome subunits, and light blue cycles represent the regulation of proteolysis. Among those, 26S proteasome non-ATPase regulatory subunit 3 homolog A (RPN3A), 20S proteasome beta subunit D1 (PBD1) and 20S proteasome alpha subunit e1 (PAE1) were enriched 100-fold in *mca-II-KOs* mutant vs WT (three-month-old seeds, adj P<0.001).
**(G)** Representative immunoblots from *MCA-II-KOc* seedling (five days-post-germination) or seeds (50 seeds/sample) with α-PAG1 or α-PBA1*.* PAG1: 20S proteasome alpha subunit G-1. PBA1: 20S Proteasome subunit beta 1. The experiment was repeated three times.
**(H)** Representative image showing etiolated seedlings grown for 5 days under darkness on mock (DMSO), 20 nM MG132, 0.3 μM concanamycin (Con A), 3.3 μM wortmannin (WM), and 5 μM E64D containing plates. Right: quantification of the hypocotyl length. The data are from three experiments, and the indicated *P* values were calculated by ordinary one-way ANOVA (*N* = 3, *n* =6-10). Note the hypocotyl inhibition caused by MG132 in WT is not seen in *mca-II-KOc* mutant, indicating that proteasome is compromised in the *mca-II-KOc* mutant.

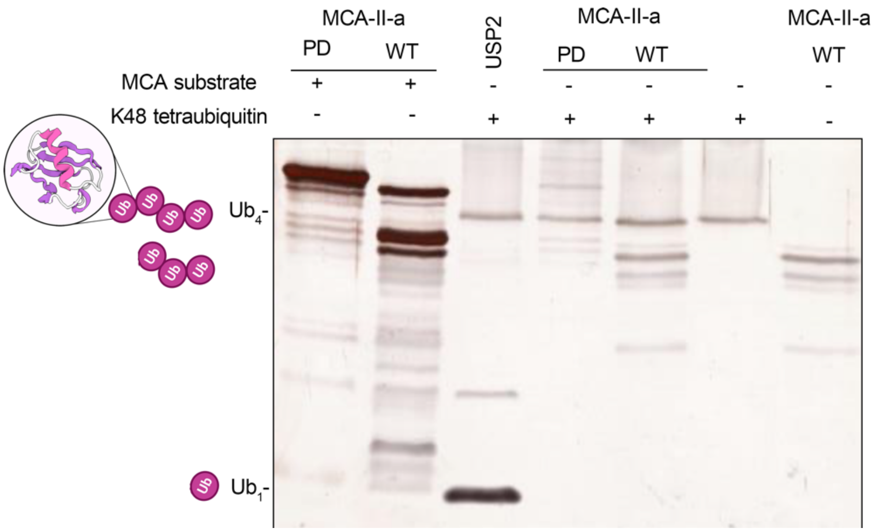


**fig. S6. MCA-IIa did not cleave K48 tetraubiquitin isopeptide linkage-type chains.** *In vitro* K48 tetra-Ub isopeptide linkage type cleavage assay was conducted using the recombinant proteins of MCA-II-a and MCA-II-a$^{PD}$ (*3, 35*). In comparison to the positive control USP2-cc (*22*) which can cleave the K48 tetra-Ub isopeptide linkages, there is no obvious cleavage catalyzed by recombinant MCA-II-a. Polyacrylamide gel electrophoresis (12%) silver-stained gel. The experiment was repeated two times.

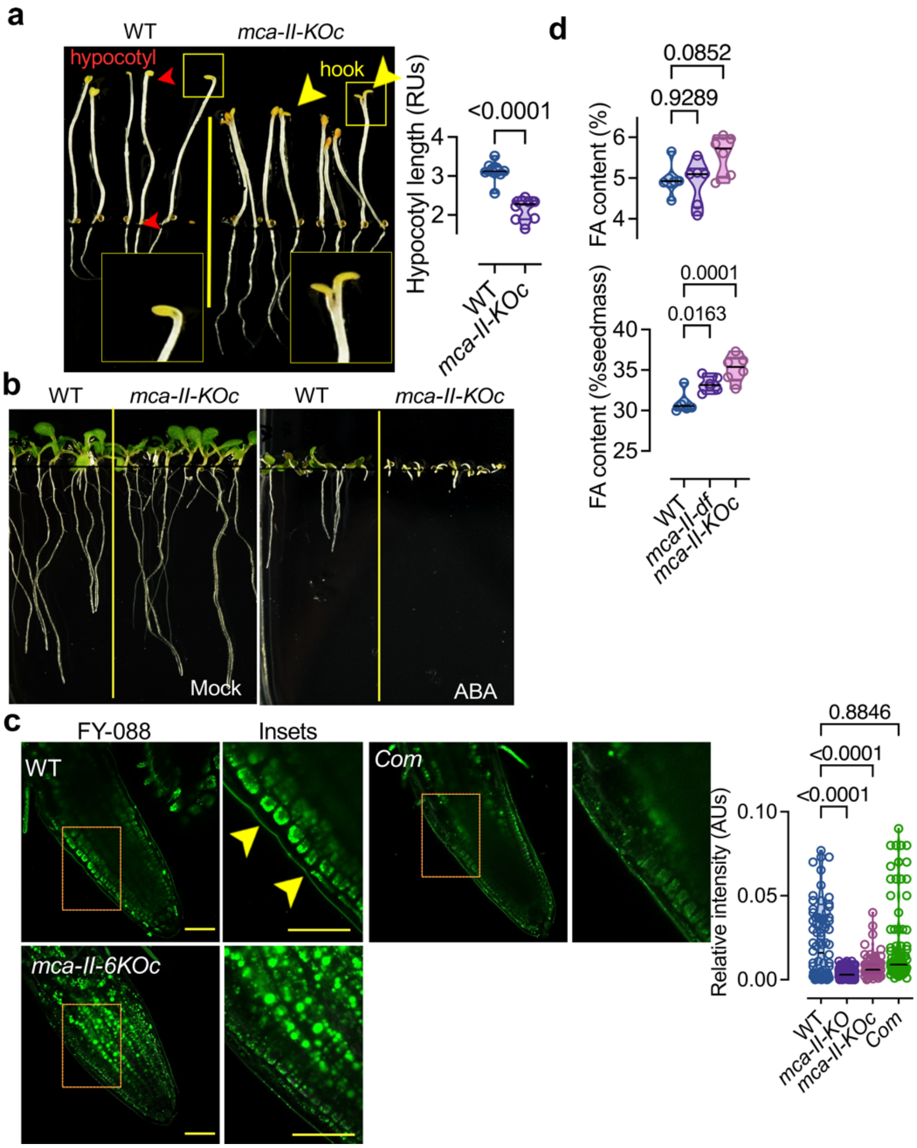


**fig. S7. The role of MCA-IIs in the regulation of PUX10 and CDC48a**

**(A)** Loss of apical hook in *mca-II-KOc* and reduced hypocotyl length. The etiolated seedlings were grown for 5 days under darkness. Red arrowheads denote the hypocotyl, while yellow arrowheads the apical hook. Right: quantification of the hypocotyl length (RU, relative units). The data are from three experiments, and *P* values were calculated by ordinary one-way ANOVA ($N$ = 3, $n$ = 33).

**(B)** Representative picture of seven-days-after-germination seedlings grown on either mock (DMSO), or ABA-containing plates (100 nM). The experiment was repeated three times ($N$ = 3, $n$ = 8).

**(C)** Representative micrograph of FY-088 staining for two-day-post-germination seedlings of WT, *mca-II-KO* (with the Cas9 cassete), *mca-II-KOc*, and a complementation line (Com, with MCA-II-a). Right: The data are from three experiments, and *P* values were calculated by ordinary one-way ANOVA ($N$ = 3, $n$ = 33).

**(D)** Fatty acid content analysis of total extracts from WT, *mca-II-KO,* and *mca-II-KOc* mutant of seeds harvested at the same time (3-month-old seeds, 100mg seeds for each line, $N$ = 5).

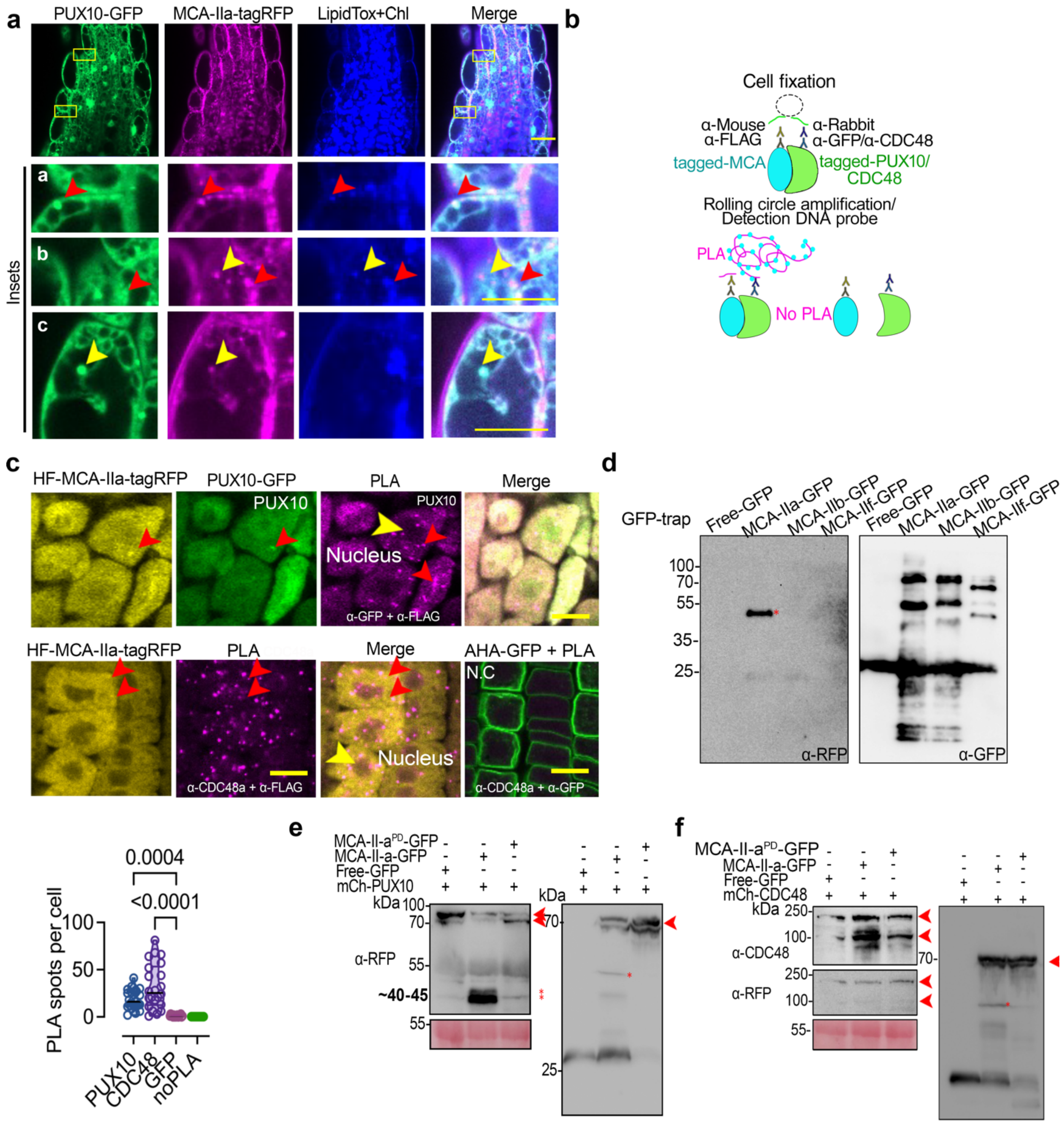


**Fig S8. MCA-II-a interaction with PUX10 and CDC48.**

**(A)** Representative confocal micrographs of LDs stained with lipidTox in the hypocotyl region of seedlings (2-DPG) of a line co-expressing *RPS5apro:MCA-II-a-tagRFP* with *PUX10pro:PUX10-GFP.* In the insets denoted with "a", red arrowheads show colocalization between MCA-II-a with PUX10; "b", red and yellow arrowheads show the MCA-II-a-tagRFP with lipidTox; "c", the signal of PUX10 or MCA-II-a did not always colocalize with LDs, especially in relatively larger droplets. Scale bar, 10 μm. The experiment was repeated multiple times.

**(B)** Cartoon showing the principle of the Proximity Ligation Assay (PLA) approach for protein-protein interaction study (< 40 nm, details in the methods) and representative confocal micrographs (lower) of PLA signal (PLA foci denoted with magenta) indicating the interaction between FLAG-MCA-II-a-tagRFP with either PUX10-GFP (α-FLAG and α-GFP) or CDC48a (α-CDC48). PLA of the P-ATPase AHA1-GFP with CDC48a is used as a negative control (α-GFP and α-CDC48). NC, negative control. Scale bars, 10 µm. Right: quantification of PLA-positive foci in cells. The data are from three experiments and the indicated *P* values were calculated by one-way ANOVA (*N* = 3, n = 8 cells/sample).

**(C)** PLA assay showing the interaction between FLAG-MCA-II-a-tagRFP with PUX10-GFP (upper) or CDC48 with FLAG-MCA-II-a-tagRFP (lower) from epidermal cells of meristematic cells from root. The experiment was done three times (N=3, n= multiple cells).

**(D)** Representative immunoblots showing coimmunoprecipitation between MCA-II-a-GFP or MCA-II-f-GFP (baits) with mCherry-PUX10 in the *N. benthamiana* transient expression system (four-days post infiltration). The red arrow indicates the proteolytic fragment of PUX10. The experiment was repeated two times (N = 2, n = 2).

**(E)** and **(F),** Representative immunoblots showing detection of mCherry-CDC48a or mCherry-PUX10 co-expressed with MCA-II-a-GFP or the inactive variant MCA-$IIa^{PD}$-GFP in the *N. benthamiana* transient expression system using α-CDC48a, α-RFP, or α-GFP. Red stars indicate the proteolytic N-terminal fragment of PUX10 (-RFP, ca. 10 kDa). The α-TUBULIN represents loading controls. The experiment was repeated two times (*N* = 2, *n* = 2).

**Other Supplementary Materials for this manuscript include the following**

**Data S1. Proteomic analyses of WT and *mca-II-KOc* 3-month-old seeds.**

**Data S2. UPR gene expression in WT or in *mca-II-KOc*.**

**Data S3. Interactions of MCA-II-a and -d obtained by TAP for WT or PD variants.**

**Table S1. Resources used in this paper.**
